# A multidisciplinary investigation into whether Andean caravans reached the southern lowlands of the Paraná-Plata basin during pre-Columbian times

**DOI:** 10.1101/2024.09.24.614829

**Authors:** Daniel Loponte, Alejandro Acosta, Tommaso Giovanardi, María J. Corriale, Owen Alexander Higgins, Mirian Carbonera, Natacha Buc, Cinzia Scaggion, Eugenio Bortolini, Giulia Marciani, Stefano Benazzi, Lucía T. Rombolá, Michael V. Westbury

**Author notes:** E-mail: Alejandro Acosta; Natacha Buc.; Tommaso Giovanardi; Cinzia Scaggion.

## Abstract

The expansion of llama caravans and the dispersal of domesticated camelids to extra-Andean regions is one of the key topics in South American archaeology, as it reflects the degree of connection between Andean societies and the surrounding lowland societies, as well as the extent of pre-Columbian pastoralism. One of the primary indicators of both processes is the presence of domesticated camelids in the archaeological record, particularly the llama (*Lama glama*). Based on somewhat ambiguous historical evidence, it has been suggested that llama caravanning may have reached the northern Pampean region and the southern Paraná River valley from an unspecified time until historical times. While the archaeological sites in these two regions have abundant camelid remains that have thus far been identified as guanaco (*Lama guanicoe*), the possibility of misidentification due to the osteometric similarity between the domestic and wild camelids could mask the presence of llamas in the record, undermining evidence of this potential major expansion of llama caravans. To clarify this issue, we applied a multidisciplinary approach combining archaeological, isotopic, and paleogenomic analyses to determine the taxonomic status of camelids recovered from archaeological sites in both areas. Our findings demonstrate that all the individuals analyzed correspond to guanacos, whose survival extended into early historical times. There is no regional evidence to support the presence of llama caravanning or domesticated camelids in the northern Pampean region or the lower Paraná River valley.

## 1. Introduction

Llama caravanning in the Andean region began during the Early Formative period (⁓ 3500 BP) and has continued into the present day. This movement of animals, goods, and people began to develop for the transportation and exchange of goods, both within the Andean area and between the highlands and the surrounding lowlands. Of particular interest in this complex mobility system are llamas (*Lama glama*), which were the pack animals primarily used in this social and economic process, with far-reaching consequences for the societies involved—consequences that can be traced in the archaeological record **[1–2]**.

Some authors **[3–4]** have suggested that llama caravanning during pre-Columbian times may have extended significantly eastward, reaching the northern Pampas plain and the lower wetland of the Paraná valley, which is adjacent to the eastern edge of this plain, in the lowlands of southeastern South America (Figure 1). According to these authors, these caravans would explain some 16^th^-century references to the presence of camelids in the Paraná Valley and the northern Pampas region, ruling out the possibility that they referred to guanacos (*Lama guanicoe*), despite the archaeological record in this region showing the existence of herds of these wild camelids. Indeed, abundant guanaco bones have been morphologically identified at Late Holocene archaeological sites of highly mobile, small bands of hunter-gatherers inhabiting the temperate northern Pampas plain, whose subsistence was based on this mammal **[5–7]**.

**Figure 1.**
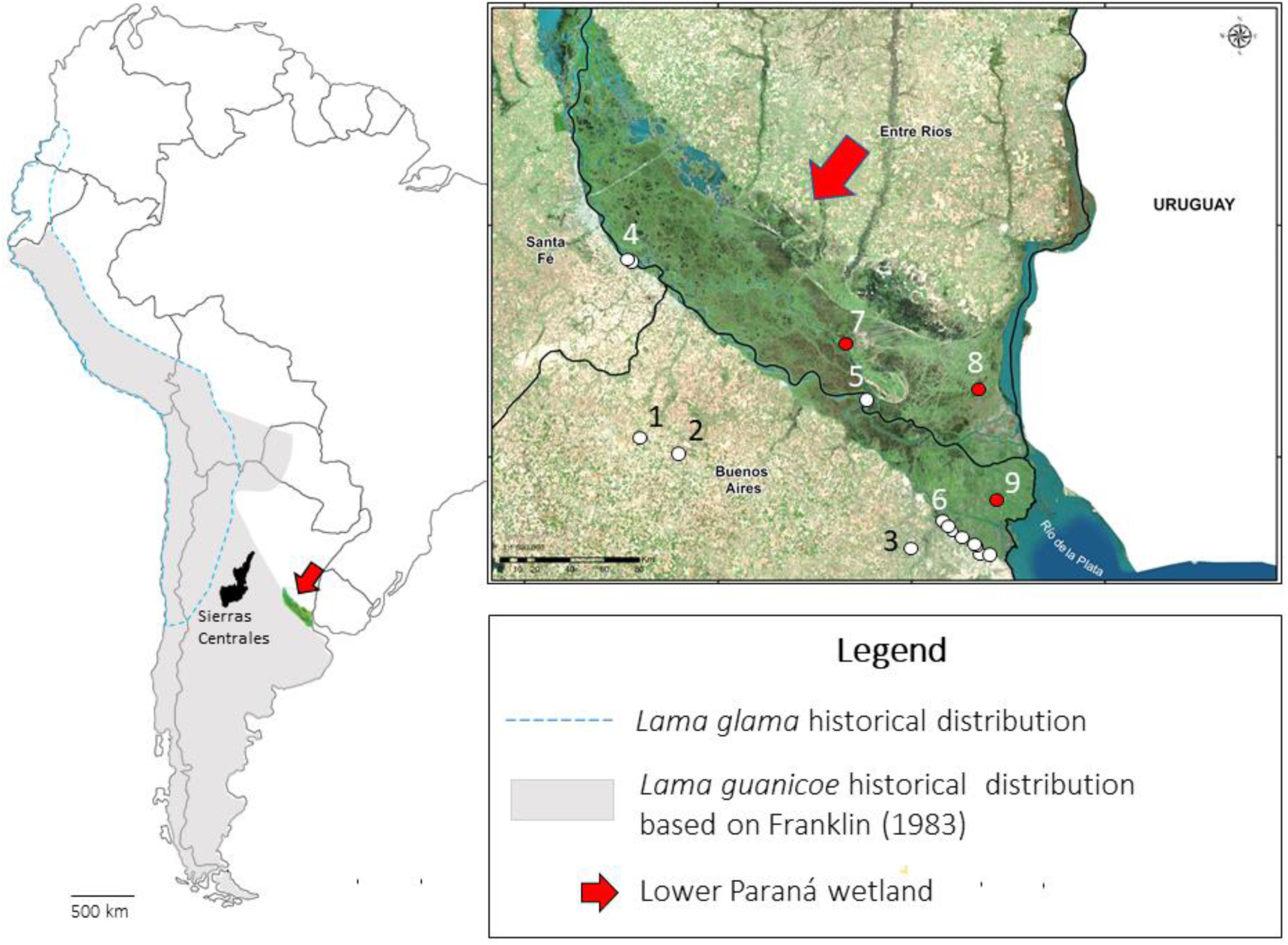
Left: Historical distribution of *Lama glama* (llama) and *Lama guanicoe* (guanaco). Redrawn from **[17]**. The Lower Paraná wetland is indicated as a green-colored area pointed with a red arrow. Right: Location of the sites included in this study. At the Pampas plain: 1 = Hunter; 2 = Meguay, and 3 = Cañada de Rocha. At the wetland on the ecotone with the Pampas: 4 = Playa Mansa and Bajada Guereño; 5 = Isla Lechiguanas 1 (near the ecotone); 6 = Túmulo de Campana site 2; La Bellaca sites 1, 2, 3; El Cazador sites 1, 2, 3; Río Luján; Garín, El Espinillo, Punta Canal; Rancho Largo; Médanos de Escobar, Guazunambí, Arroyo Sarandí. Sites deeply inside the wetland: 7 = La Argentina. 8 = Cerro Lutz and Las Ánimas. 9 = Arroyo Fredes.

There is also a fragmented and sparse record of guanaco bones at Late Holocene archaeological sites in the adjacent wetlands of the lower Paraná valley, associated with more sedentary, complex hunter-gatherers whose economy relied on fishing and the exploitation of mammals typical of this subtropical wetland, alongside ancillary horticulture **[8]**. In these sites, guanaco bones are primarily represented by metapodials and phalanges, which has been interpreted as the result of logistical hunting expeditions into the adjacent northern Pampas plain to obtain guanaco hides, or hide trade between the hunters of the northern Pampas plain and those inhabiting the wetland **[9]**. This interpretation is based on the fact that guanacos were not living in the swampy environment of the Lower Paraná wetland. Guanaco hides are of significantly higher quality than those of all other fur-bearing animals from the Pampean plain and the Paraná valley, as they possess several properties unique to the region: they are very flexible, thick, have a high hair density, and are large in size. This likely motivated the indigenous groups inhabiting the wetland to seek guanaco hides from the adjacent plain. The archaeological evidence for the introduction of hides into ecotone sites includes the presence of phalanges and distal metapodials, with the latter often fractured at their metaphyses. These anatomical elements frequently remain attached to hides after skinning, classifying them as “riders” in ethnoarchaeological and archaeological studies **[10]**. The deliberate fragmentation of the metapodials appears to be a strategy for removing the hide attached to the bone and reducing the overall weight of the hides, even though the weight of these bones is negligible compared to that of a freshly skinned guanaco hide.

If llama caravanning from the Andean region had reached these lowlands, they would have left evidence in the region, including the presence of llamas which could be present in the archaeological record of both the northern Pampean plain and the Paraná valley, mistakenly identified as guanacos. While it is highly unlikely that the highly mobile hunter-gatherers of the northern Pampas plain “hunted” llamas or consumed llamas provided by these supposed Andean caravans as part of their subsistence, the case could be different for the more complex societies of the Paraná valley, making this an intriguing point to explore due to the enormous implications that the identification of pre-Columbian llamas in southeastern South America would have. Additionally, this identification would also support the existence of llama caravanning or the presence of llamas in intermediate areas between the Andean region and the Paraná valley, such as the Sierras Centrales Hills (Figure 1), where its positive identification (based on osteometry) has so far been elusive **[11]**.

To address the issues outlined above, we reassess the taxonomic status of camelids from archaeological sites in the northern Pampas region and the lower Paraná valley (Figure 1). Given the well-known difficulty in osteometrically distinguishing llamas from guanacos (**[12–16]**, and the references therein], and the limitations this method imposes on our objectives (see Supplementary Text), we applied a complimentary set of analytical approaches. We first examined the anatomical representation and age classes of camelids in both areas in order to identify the human behaviors that led to the creation of the archaeological record of these camelids in both regions. Secondly, we used isotopic provenance markers (δ^18^O, ^87^Sr/^86^Sr) to assess whether the camelids from both areas are local organisms and whether there is a correspondence between the guanacos from the Pampean plain and those camelids recovered from sites in the Paraná valley. Third, we employed a dietary marker (δ^13^C) to explore whether there were differences in the diets of guanacos from the Pampas plain and camelids from the ecotonal sites, which would be expected if they represented wild and domesticated populations. Lastly, we compared the genomes of camelids recovered from both the Pampas plain and the ecotonal sites of the wetland in the lower Paraná valley with a modern reference panel of all extant South American camelids (sub)species. For further information on archaeological contexts, sampling strategies and taphonomic biases, see the Supplementary Text.

## 2. Materials and Methods

### 2.1. Archaeological and isotopic analysis

For this study, we analyzed the bones of camelids recovered from three sites located in the northern Pampas plain (Meguay, Hunter and Cañada de Rocha) and from 21 sites located at the wetland in the ecotone with the Pampas plain (Figure 1 and Supplementary Tables 1 and 2). Values of δ^18^O and δ^13^C were obtained at the Environmental Isotope Laboratory, Department of Geosciences of Arizona University, and Instituto de Geocronología y Geología Isotópica (CONICET - University of Buenos Aires). A few samples were analyzed at the Center for Applied Isotope Studies (CAIS, University of Georgia). Protocols and methods are described in the Supplementary Text. Values of ^87^Sr/^86^Sr were obtained from second set of samples at the Centro Interdipartimentale Grandi Strumenti (CIGS) of the University of Modena and Reggio Emilia (Italy). Protocols and methodologies are described in Supplementary Text. Statistical analysis of the obtained isotope data was performed using general linear mixed models (GLMM). These response variables were fitted to a model assuming a normal distribution, with site and species considered as random factors. *Post-hoc* mean comparisons were conducted using a Tukey test. To evaluate differences between species, the site was included as a random factor. We assessed distribution adjustment graphically from complete model residuals. In cases of heteroscedasticity, this was modeled using the varIdent function. All analyses were performed using R version 4.2.2 **[18]**, utilizing lmer function from the ‘lme4’ package **[19]** and lme function from the ‘nlme’ package **[20]**. All tests were two-tailed, and differences were considered significant at p < 0.05.

### 2.3. DNA

#### 2.3.1. Data generation

We obtained between 50-100mg bone powder from each sample using a Dremel drill piece. DNA was extracted from the bone powder and converted into Illumina sequencing libraries using a TECAN Fluent780 laboratory robot at the University of Copenhagen. Bone powder was first demineralized using an extraction buffer pre-digestion step for 30 minutes as described previously **[21]**. Following pre-digestion, DNA extractions were performed by combining 150 µl of demineralized material with 1.5 ml binding buffer (500 ml Qiagen PB, supplemented with 15 ml Sodium acetate 3M, and 1.25 ml 5M NaCl, phenol red, adjusted to pH = 5) and 10 µl of paramagnetic beads for 15 minutes **[22]**. Pelleted beads were washed twice in 450 µl and 100 µl 80% ethanol + 20% 10mM Tris-HCl, respectively, and eluted in 35µl of 10 mM Tris-HCl + 0.05% Tween-20. USER treatment was performed for each extract by adding 2.5 µl USER enzyme and 7.5µl water followed by an incubation at 37°C for 3 hours. Double-stranded Illumina libraries were prepared following the protocol described by **[23]**. Clean-up procedures after end-repair and adapter-ligation were performed with 10 µl of paramagnetic beads in a 10x volume of the binding buffer described above. The number of PCR cycles required was evaluated by qPCR using 1µl of pre-amplified library. Indexing PCR was performed using 8-bp unique dual indexing with KAPA HiFi HotStart Uracil+ according to manufacturer’s recommendations. Final purification of libraries was performed using a 1:1.6 ratio of library to HighPrep™ PCR beads. Length distribution and concentration of individual purified libraries was controlled using the Fragment Analyzer (High Sensitivity kit). Amplified sequencing libraries were sent to Novogene (UK) for sequencing of ∼5Gb on an Illumina Novoseq sequencing platform using paired-end 150bp read chemistry. To generate a reference dataset for subspecies level genomic identification we downloaded the raw sequencing reads from 41 South American camelids, with representatives from all four extant species. For further population identification, we downloaded raw reads from an additional 34 *L. g. guanicoe* individuals.

#### 2.3.2. Data processing

We removed Illumina adapter sequences, low-quality reads (mean q <25), short reads (<30bp), and merged overlapping read pairs with Fastp v0.23.2 **[24]**. We mapped the trimmed reads to the alpaca (*Vicugna pacos*) reference genome Genbank accession: GCF_000164845.3) using Burrows-wheeler-aligner (BWA) v0.7.15 **[25]**, and either utilising the aln algorithm, with the seed disabled (-l 999) (otherwise default parameters) for the ancient individuals or the mem algorithm with default parameters for the modern individuals. We parsed the alignment files and removed duplicates and reads of mapping quality score <30 using SAMtools v1.6 **[26]**. We checked for ancient DNA damage patterns of the ancient individuals using mapdamage2 **[27]**.

We identified scaffolds putatively originating from the sex chromosomes in the alpaca assembly by aligning the assembly to the Cow (*Bos tauraus*) X (Genbank accession: CM008168.2) and Human (*Homo sapiens*) Y (Genbank accession: NC_000024.10) chromosomes using satsuma synteny v2.1 **[28]** with default parameters. We followed the SeXY pipeline **[29]** to identify the sex of each of the ancient individuals. We set a minimum threshold of 5,000 mapped reads to ensure the reliability of the results. If an individual had an X:A ratio of <0.7 it was designated as a male. If an individual had an X:A ratio of >0.8 it was designated a female. Individuals with ratios between 0.7 and 0.8 were deemed undetermined.

#### 2.3.3. Species identification

To identify the potential (sub)species of the archaeological individuals we used four different methods: Principal component analysis (PCA), admixture proportions, phylogenetic trees, and pairwise distances. We only included individuals with >1.2Mb of data (>0.0006x average genome-wide coverage). We performed three different PCAs, one using all extant South American camelid subspecies, one using only those from the *Lama* genus, and one using only those from the *L.g.guanicoe* subspecies. As input, we computed genotype likelihoods (GL) with ANGSDv0.921 **[30]** using only the autosomal scaffolds >10Mb in length and the following parameters: -minmapQ 20 -minQ 30 -GL 2 -doGlf 2 -doMajorMinor 1 -rmtrans 1 -doMaf 2 -SNP_pval 1e-6 -minmaf 0.05 -skiptriallelic 1 -uniqueonly 1. In both cases, we set a minimum individual threshold for the number of modern individuals +1, i.e. 42 for the complete dataset, 18 for the *Lama* only dataset, and 41 for the *L.g.guanicoe* dataset. We converted our GL into a covariance matrix using PCAngsdv0.98 **[31]**.

To calculate the admixture proportions, we used the same GL as input to PCAngsd but with the additional parameter of -admix. The most probable K was selected by PCAngsd based on Velicier’s minimum average partial (MAP) test.

We computed a distance matrix for all individuals using a pseudohaploid consensus base call approach in ANGSD. We used the same filtering parameters as for the GL but with the additional parameters -doIBS 2 -minminor 2 and -makematrix 1. We built an unrooted neighbour joining phylogenetic tree from the output distance matrix using fastme v2.1.6.1 **[32]** and default parameters.

We also computed pairwise distances between the modern *Lama* specimens and the ancient camelids using the same consensus base call identity by state approach in ANGSD as used for the *phylogenetic* tree. We extracted only comparisons that included our ancient individuals against the modern individuals.

## 3. Results

### 3.1. The anatomical representation and chronology of camelids in the northern Pampean plain and the Lower Paraná Valley

In the three excavated sites on the northern Pampas plain (Cañada de Rocha, Hunter, and Meguay), located 40 to 80 km from the western boundary of the ecotone with the wetland (Figure 1), camelid bones dominate the archaeofaunal assemblages, with smaller quantities of armadillos (e.g., *Chaetophractus villosus; Zaedyus pichiy*), pampas deer (*Ozotoceros bezoaticus*), and rhea (*Rhea americana*) remains. The preservation of the faunal record is not influenced by bone density or biased by culinary or religious practices (Supplementary Text). The anatomical representation of camelids is complete, including skulls, ribs, scapulae, pelvises, forelimbs, and hindlimbs. The long bones of the camelids show fresh fractures, suggesting the consumption of marrow (**[5,6]**; see also Supplementary Text). Associated artifacts are bola stones intended for hunting large prey such as guanacos, deer and rhea, as well as a few quantities of pottery and lithic tools typical of local hunter -gatherers of the Pampas plain. These lithic tools were made from rocks obtained in the southern Pampean hills (quartzites and highly silicified translucent microcrystalline rocks). None of these sites present non-local taxa (e.g., Andean species), Andean raw materials, metals, or Andean pottery, nor dung deposits or corral structures **[5–6]**.

Two AMS radiocarbon dates obtained from camelid bones recovered from the Hunter and Meguay sites date between 51 BCE and 196 CE, and between 897 and 1022 CE, respectively. Five AMS radiocarbon dates from the Cañada de Rocha site yielded ages ranging between 1324 and 1615 CE (Supplementary Table 1 and the references therein). Other radiocarbon ages were also obtained from the latter site, but the samples were altered or contaminated according to (**[33]**, p. 189-192). All dated samples from these sites in the northern Pampean plain underwent an Acid-Base-Acid treatment to remove any contaminants **[34]**. All the samples yielded sufficient collagen (> 1%) and carbon mass (> 1 mg) for AMS dating, which is a more reliable method than conventional ^14^C dating due to a lower likelihood of contamination and greater accuracy. In two of these five samples, the C/N ratio was also measured, yielding values at 3.23 and 3.24 **[7]**, demonstrating the absence of diagenetic evidence and confirming the reliability of the radiocarbon ages obtained. Moreover, these two dates largely overlap with the other three obtained from guanaco bones from the same site, which, combined with the analytical quality of the dated samples, makes the chronology of these five guanaco specimens extremely robust, without outliers. Therefore, these dates obtained directly from guanaco bones, demonstrate that these animals inhabited the northern Pampean plain during the Late Holocene and reaching the onset of the historical period, with a range extending into the early post-Columbian period, when the first descriptions of the region were made, documenting the presence of camelids in the area (Supplementary Table 1 and Supplementary Figure 1).

From the wetland sites, 21 archaeofaunal records with camelids bones were analyzed by several authors, which are mostly composed of fish bones (Supplementary Text). The mammal bones correspond to taxa with habitats within the wetland, such as the marsh deer (*Blastocerus dichotomus*) and coypu (*Myocastor coypus*). These sites also contain a small quantity of bones from species typical of the adjacent Pampas plain, such as pampas deer, rhea, and bones that have been assigned to *L. guanicoe* but which we will, for now, identify only as Camelidae sp. in order to assess whether they belong to *L. guanicoe* or *L. glama.* The bones of camelids are found in quantities lower than 0.1 %NISP, while those of *O. bezoarticus* can reach up to 5 %NISP (Supplementary Text). This is likely due to the fact that the latter species has a more flexible behavior compared to guanaco, allowing it to inhabit swampy environments fragmented by bodies of water and vegetation. The bone preservation in the wetland sites is not influenced by culinary practices, bone mineral density or religious practices (Supplemental Text).

Several camelids bones from these sites were directly dated and others were recovered in primary association with dated samples ranging from ⁓ 700 BCE to 1627 CE (Supplementary Table 2). Among the large number of camelid radiocarbon ages and other associated taxa from the wetland sites, it is worth highlighting the one from the Playa Mansa site (Supplementary Table 2). Here, a guanaco bone was directly dated using AMS, with an age range between 719 BCE and 233 BCE (95.4%) or between 544 and 378 BCE (92.8%). Either of these ranges predates the appearance of llamas in the microthermal valleys and surrounding lowlands of Northwest Argentina **[15]**. Therefore, it is unlikely that this bone corresponds to a llama. Furthermore, it indicates that the human behavior responsible for this peculiar record, concentrated in phalanges and metapodials at the Paraná valley sites, was active before the expansion of llamas into those valleys of the Northwest Argentina, and that this same mechanism remained active until the 16^th^ century (see Supplementary Table 2).

Despite the fact that the ecotone sites containing camelid bones correspond to the same period as the adjacent sites in the Pampean plain, the anatomical representation of camelids is completely different, as only metapodials and phalanges were predominantly recovered at the former sites (Supplementary Table 2). In just two sites two patellae (one in each) and one tooth were recovered (Supplementary Table 2). Therefore, the significant differences in the anatomical representation of camelids correlate with the type of environment, suggesting that this representation is influenced by environmental factors. In the wetland sites, the almost exclusive presence of metapodials and phalanges indicates a specific behavior, possibly related to the procurement of guanaco hides from the adjacent plain and certain by-products that would have increased the return of logistical hunting expeditions, such as guanaco leg tendons or the use of metapodials as blanks for manufacturing long and highly resistant bone points **[35]**. At none of the ecotone sites with camelid remains were pottery, art, or rocks of Andean origin recovered. Nor were any camps attributable to llama caravans, Andean tombs, or Andean animal species identified in the area.

### 3.2. Age classes

The metapodials of guanaco fuse at approximately 30 months of age **[36]**. At the Hunter and Meguay sites located in the northern Pampean plain, unfused metapodials account for around or less than 10% of the total recovered. The emphasis in capturing animals here appears to have been focused on acquiring adult individuals, as they provide the largest biomass, although some juveniles were captured in smaller proportions. In contrast, at the ecotonal sites, of the 43 metapodial condyles included in this study, 29 are fused with their respective metaphyses (∼67%), while the remaining 14 condyles (∼33%) are unfused (Supplementary Table 3). The greater selectivity for capturing juvenile animals at the wetland ecotone sites strongly supports the hypothesis of logistic guanaco hunting (or acquisition through exchange) specifically aimed at obtaining hides, as juvenile skins are more flexible, resistant, and of better quality than those of adults **[37]**. This also aligns with the osteological record, which focuses primarily on metapodials and phalanges previously discussed (see Section 1). Additionally, it is worth noting that, of the six adult animals genetically sexed at these wetland ecotone sites, five were females and only one was a male. Although the possibility of sampling bias cannot be ruled out, the combination of a higher proportion of females and the significant presence of juveniles suggests that the captures may have focused more on family groups of guanacos, where there is a greater predominance of females and juveniles up to approximately one year of age **[38]**. However, to confirm this hypothesis, larger sample sizes will be necessary.

3.3. δ^18^O

We obtained 28 δ^18^O values from different species typical of the Paraná wetland (subset 1), which were recovered from various sites located in the ecotone (Supplementary Table 4). These values average −3.24 ± 0.7 ‰, consistent with local organisms from the Paraná Valley (**[39]**; see also Figure 2-A and Supplementary Text). A second subset of samples from these ecotone sites, comprising five specimens from Pampean taxa (four pampas deer and one rhea), yielded an average of −0.64 ± 1.18 ‰. Meanwhile, a third subset, consisting of eleven camelid samples also recovered from various ecotonal sites, yielded an average of −0.76 ± 0.7 ‰. These latter two subsets (2 and 3) exhibit values that align with those of organisms with life cycles in the adjacent Pampas plain (see Supplementary Text) and show no statistically significant differences between them (F = 2.06; p = 0.179), allowing us to treat them as a single group (hereafter referred to as the “combined group”). When comparing this combined group (subsets 2 + 3 = 16 samples) with subset 1 (n = 12, wetland species), we find a statistically significant difference (F = 25.99; p = 0.003). This difference remains statistically significant when focusing solely on the mammals in the combined group (subsets 2 + 3) compared to mammals in subset 1 (F = 64.92; p < 0.001). These results indicate that the faunal assemblages recovered from the wetland sites comprise organisms with diverse life histories. Some developed their life-cycles in the wetland (subset 1), while others (combined group; subsets 2 and 3) inhabited the adjacent northern Pampas plain. This demonstrates that Pampean taxa, such as deer and camelids, coexisted in the same environment and that hunter-gatherers with campsites in the wetland ecotone exploited or acquired resources from the adjacent plain, including pampas deer, rhea, and camelids.

**Figure 2.**
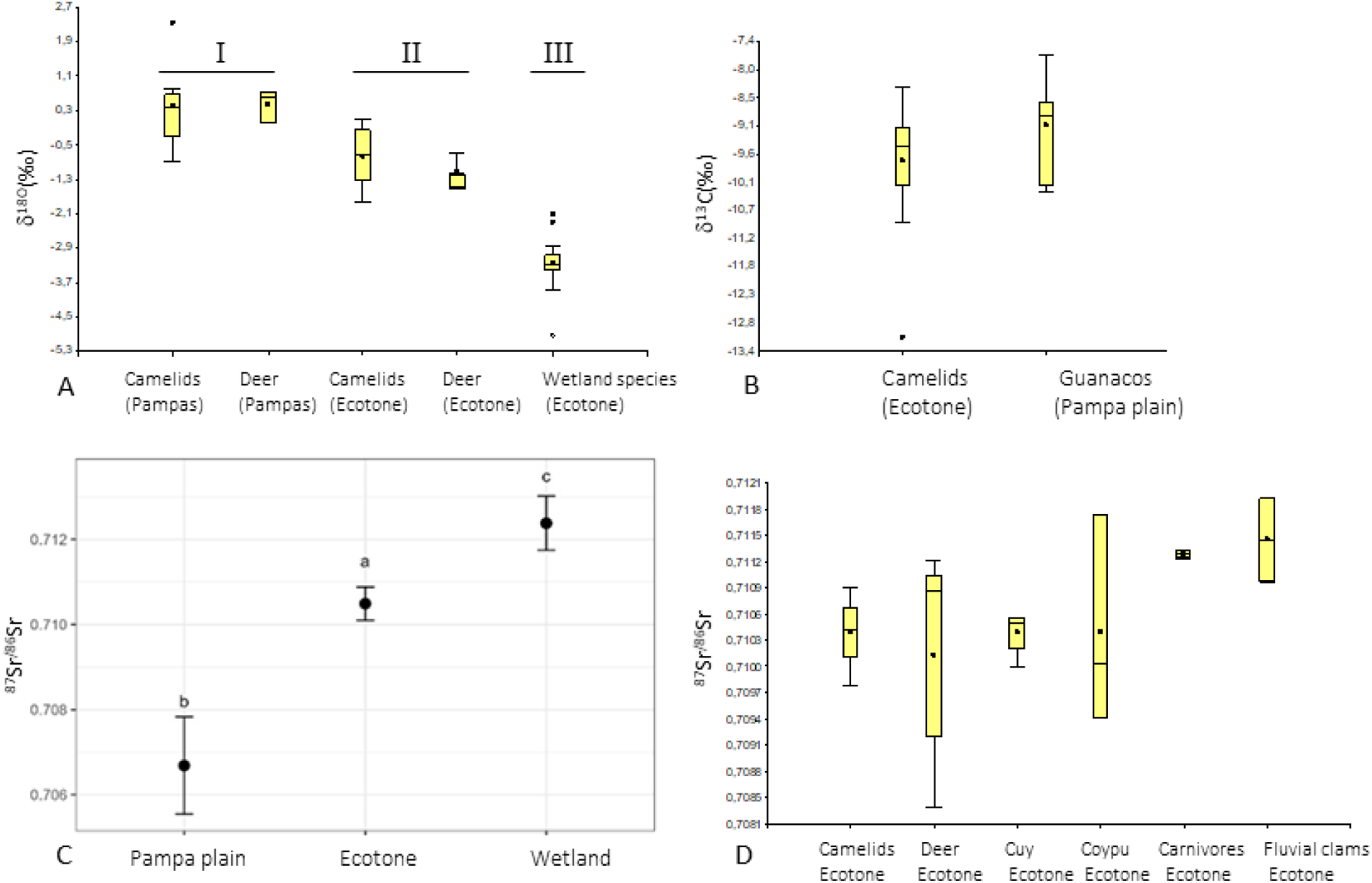
**A** = δ^18^O values of samples grouped by taxa and provenience area, except for wetland species which are clustered in a single group (significant statistical differences between groups I, II and III; see text). **B** = δ^13^C values for camelids recovered at the ecotonal sites in the wetland and Pampas plain (no significant statistical differences). **C** = Confidence intervals of ^87^Sr/^86^Sr values according to the area of origin of the samples (all taxa included in each area) with significant differences between them. **D** = ^87^Sr/^86^Sr values of different taxonomic groups based on samples recovered from the ecotonal sites located at the wetland (no statistical differences between them).

At the same time, the combined group (subsets 2+3) shows significant differences from the twelve samples recovered from sites located in the deep interior of the Pampas plain (Hunter and Meguay sites), consisting of eight camelids, three pampas deer and one rhea (*F* = −3.8372*; p* < 0.001) (see raw values in Supplementary Table 4). These differences persist when comparing only the mammals (*F* = 20.57; *p* < 0.001), camelids (*F* = 9.72*; p* = 0.0062) or deer *(F* = 33,62; *p* = 0.002). These differences are primarily due to the fact that the δ^18^O values of Pampean taxa recovered from wetland sites show, on average, slightly more negative values than organisms that lived deeper within the plain (Figure 2-A). This is expected given the environmental effect of the Paraná River over an approximate area of 10 km towards the interior of the plain, due to the ambient humidity it provides and the inflow of Paraná water into the estuaries of the rivers descending from the interior of the north Pampean plain **[39, 40]**. All these results suggest that camelids and deer (*O. bezoarticus*) recovered from wetland sites underwent their life-cycles in the northern Pampean plain near the ecotone where they hunted.

### 3.4. δ^13^C (inorganic fraction)

The δ^13^C (apatite) values of twelve camelid samples recovered from the wetland ecotone sites average −9.73 ± 1.2 ‰ (Supplementary Table 5). This mean remains practically unchanged if the outlier from the Garín site is removed, whose δ^13^C value (−13.1 ‰) is practically monoisotopic C_3_. Considering that pure C_3_ diets yield values in the inorganic fraction as low as −13.5 ‰, and pure C_4_ diets ∼ −3 ‰ **[41–44]** the diet of these camelids was predominantly C_3_ with a smaller component of C_4_ plants. The coefficient of variation of these twelve samples is approximately 10 %, indicating a homogeneous diet typical of organisms not affected by human interference and different from the mostly enriched and highly variable values found in llamas **[45–48]**. On the other hand, eight guanaco samples from sites located at the northern Pampas plain have similar mean and dispersion values: −9.03 ± 0.87 ‰. Both sets do not exhibit statistically significant differences (*F* = 1.37*; p* = 0.12), demonstrating a similar total diet (carbohydrates + lipids + proteins) (Figure 2-B). Although there are no statistically significant differences between the camelids from the interior of the Pampas plain and those from the wetland, there appears to be a slight tendency toward higher δ^13^C values in the former. This is expected, as environmental humidity decreases to the west due to the reduced humidifying effect of the Paraná River, leading to increased aridity and a greater prevalence of C_4_ grasses **[40, 49, 50]**. On the contrary, the more negative values for the camelids recovered in the wetland ecotone perfectly fit a more humid environment of the Pampean plain near the ecotone with the wetland, with a greater availability of C_3_ grasses and shrubs (also C_3_), as well as a larger supply of trees for browsing (C_3_) than in the deep interior of the Pampas plain, which lacked trees **[41]**.

### 3.5. ^87^Sr/^86^Sr

We obtained 55 ^87^Sr/^86^Sr values for this study (Supplementary Table 6). The samples from the Pampas plain include six guanacos and two smaller mammals averaging 0.706684 ± 0.000028. No differences up to the fourth decimal, and very small differences in the fifth decimal, which reflects the monotonous geology of the northern Pampean plain where the Hunter and Meguay sites are located (Supplementary Table 6 and Supplementary Text).

The samples from the wetland ecotone sites were divided into two subsets. The first one includes 14 samples of non-wetland taxa such as pampas deer (*n* = 4), rhea (*n* = 1), and camelids (*n* = 9) which gave an ^87^Sr/^86^Sr mean of 0.710259 ± 0.000784. Within this subgroup, camelids and pampas deer show no statistically significant differences among them (*F* = 0.0026; *p* = 0.96), echoing the same environment for their development and life cycles. The second subgroup from the wetland ecotone sites is composed of twelve samples of typical wetland taxa, including two carnivores, small to medium-sized rodents that are practically sedentary (C*avia aperea* and *Myocastor coypus*), and mollusks (*Diplodon* sp.) with even more localized life cycles, localized in this case at the ecotone of the wetland. This second subset has an mean value of 0.710654 ± 0.000728. This mean is identical to the third decimal place compared to the camelids and deer recovered at the wetland ecotone sites. In fact, both subgroups of the ecotone sites yielded nonsignificant differences (*F* = 2.01; *p* = 0.24). Moreover, no differences are observed when comparisons are made considering taxonomic categories either (Figure 2-D).

The similarity in the ^87^Sr/^86^Sr values between taxa typical of the Pampas plain and those from the wetland, as recovered in the ecotonal sites, stems from the fact that they completed their life cycles in the ecotone and surrounding areas, where sediments from each geomorphological unit intersect (Supplementary Text). As a result, local aorganisms living in the ecotone reflect an averaged value from these intersecting formations, producing characteristic values for the region, which allows them to be grouped as typical of this ecotonal area. These results redundantly demonstrate that the camelid remains recovered from the ecotone sites are local resources.

The third group anayzed corresponds to 21 samples of sedentary species (*C. aperea*, *M. coypus*, *H. hydrochaeris* and *Diplodon* sp.) and four fishes from sites deep inside the wetland and intermediate position (Figure 1), with mean values of 0.712385 ± 0.00107. These three major groups (deep interior of the Pampas plain, ecotone, and deep interior of the wetland, including two intermediate samples) show statistically significant differences between them (*F* = 70.0; *p* < 0.001) (Figure 2-C). This is highly relevant as it allows us to categorize the areas of origin of the faunal resources in relation to these three main landscape units.

### 3.6. DNA

#### 3.6.1. Mapping

After mapping to the alpaca reference genome, all modern samples had relatively high genome-wide coverages ranging from 8.34x to 41.11x (Supplementary Table 7). As for the ancient samples, coverages ranged between ∼0.000 and 0.0017x or 2.5 kb mapped base pairs and 3.5Mb (Supplementary Table 7). The ancient samples showed typical ancient DNA damage patterns with elevated levels of A-G and C-T transitions at the read ends (Supplementary Figure 2) and short read lengths of ∼50bp (Supplementary Table 8).

#### 3.6.2. Sex determination

Using the SeXY pipeline we identified six females and two males (Supplementary Table 8). The sex of 14 individuals could not be identified due to low numbers of mapped reads (<5,000).

#### 3.6.3. Species identification

The PCA (Figure 3A-C), pairwise distances (Figure 3-D), admixture proportions (Figure 3-E), and phylogenetic tree (Supplementary Figure 3) all showed the Pampas plain and ecotone individuals to be more closely related to the *Lama* genus than to *Vicugna.* Within *Lama*, all analyses suggested a closer relationship between the ancient individuals and *L. g. guanicoe* relative to *L. g. cacsilensis / L. glama*. In the *Lama* only PCA analysis (Figure 3-B), the archeological samples fall into the *L. g. guanicoe* cluster. In the *L. g. guanicoe* only PCA, the ancient individuals fall closest to but not within the individuals from a similar latitude. This could suggest unique ancestry not present in living populations (Figure 3-C). The admixture proportions show that their ancestry mainly consists of the same as the modern *L. g. guanicoe* individuals. Two *L. g. cacsilensis* (Cacsilensis1 and Cacsilensis2) individuals also fall into the *L. g. guanicoe* cluster but the study in which this data was originally published suggested that these two may have actually been mislabelled guanaco **[51]** so we adjusted our results while taking this into account.

**Figure 3:**
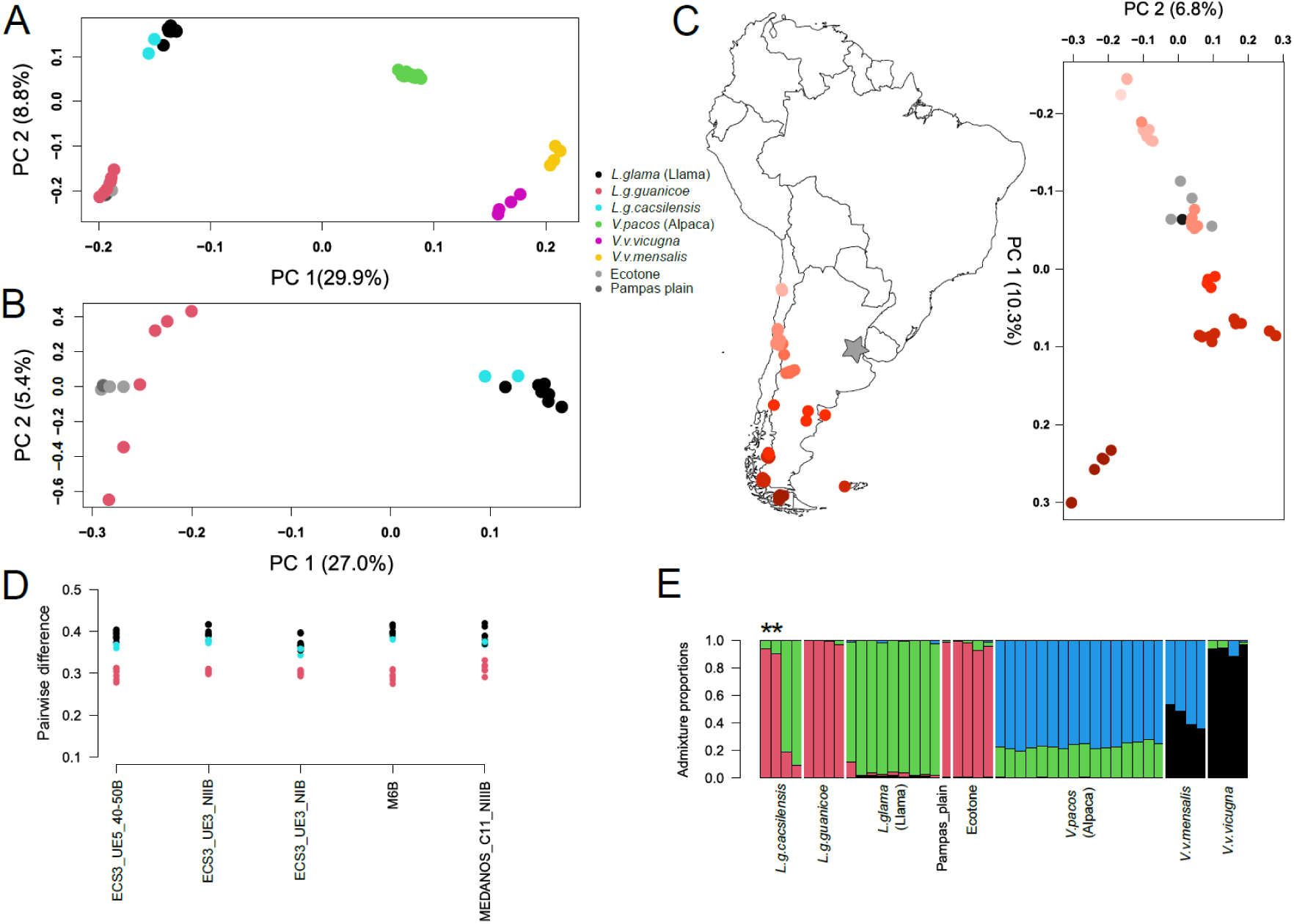
Genomic analysis of modern camelid individuals with the ecotone and Pampas plain specimens. PCA using a reference panel containing either (A) all modern individuals, or (B) just *Lama individuals,* or (C) just *L.g.guanicoe* individuals. Insert map shows the colours of the contemporary *L.g.guanicoe* in the PCA. (D) Pairwise distance comparisons calculated using only the *Lama* reference panel in ANGSD. (E) Admixture proportions taken from the GL using a K of 4 as determined by PCAngsd and the entire modern individual reference panel. *Denotes the two potentially mislabeled *L. g. cacsilensis* that cluster with *L. g. guanicoe* individuals.

The phylogenetic tree and pairwise distances also support an overall closer relationship to *L. g. guanicoe* with the branches of the wetland being closer to *L. g. guanicoe* in the neighbour joining tree (Supplementary Figure 3) and the pairwise distances being slightly lower (Figure 3-D).

## 4. Discussion

Our results clearly identify the camelid bones from the northern Pampean plain and wetland ecotone sites as *Lama guanicoe guanicoe*, with no evidence indicating the presence of domesticated camelids. AMS dating conducted on bone samples from these guanacos, which have appropriate preservation and quality analytical parameters, demonstrates that they persisted into the historical period. This provided early European travelers the opportunity to observe and describe them under various names, including “Peruvian sheep,” which led to the erroneous belief that they were domesticated camelids. **[9]**. Late Holocene pre-Columbian hunter-gatherers in the northern Pampean plain hunted guanacos, which formed the basis of their subsistence. These hunts took place near their residential camps, which explains the homogeneity of the ^87^Sr/^86^Sr and δ^18^O values, as well as the complete anatomical representation of these large animals in the archaeological sites of the plain. Since most of the hunted animals were adults, the selection of these hunters was biased towards individuals that provided the greatest amount of food, while juveniles made up a smaller fraction. The guanacos from these northern Pampean plain sites, which we analyzed here, show isotopic values reflecting the grazing areas of the northern Pampean plain, with a higher proportion of C_3_ grasses and a smaller fraction of C_4_ grasses. All this evidence demonstrates that these animals were local, and that the northern Pampean plain, adjacent to the Paraná River valley and its wetland, supported guanaco herds until historical times, serving as a source of food and raw materials for the indigenous populations of the area.

At the wetland ecotone sites, guanaco remains are represented almost exclusively by phalanges and metapodials, along with a high proportion of juveniles. This pattern clearly suggests that these animals were not hunted or acquired for food, but rather for their hides. The ^87^Sr/^86^Sr and δ^18^O values indicate that these guanacos were local to the Pampean plain, adjacent to the wetland ecotone. Their diet is indistinguishable from that of the guanacos recovered from sites in the northern Pampean plain, which is consistent with the development of a largely homogeneous steppe with minimal clinal variations in vegetation. The isotopic data further rule out the possibility that these were non-local animals or that they had been subjected to selective feeding or grazing practices.

The notable presence of juvenile guanacos at the wetland ecotone sites points to a specific interest in hunting or acquiring young animals, as juvenile hides are more flexible, making them easier to process and better suited for producing higher-quality blankets, bags, and ropes **[37]**. It is also worth noting the skewed sex ratio among adult guanacos in these sites, with females outnumbering males by a ratio of 5 to 1 (Supplementary Table 8). While this may be the result of sampling bias, it could also reflect hunting practices focused on family groups, which tend to have a higher proportion of females and juveniles under one year old **[37,38, 52]**.

The practice of guanaco hunting from the wetland ecotone sites, or their acquisition through trade with groups from the northern Pampas plain, was carried out continuously from at least ⁓700 BCE, which corresponds to a period prior to the expansion of llamas into the lower valleys of northwest Argentina. It is likely that this logistical hunting or trade practice is even older, as it has already been identified at the Playa Mansa site, which is the oldest ecotone site identified so far from these complex societies of the lower Paraná valley. This same pattern extends to later sites, demonstrating the continuity of this behavior until historical times.

Overall, the results indicate that there is no archaeological evidence of llamas in the northern Pampean plain or in the wetland ecotone of the lower Paraná River. Instead, there were herds of guanacos in this region until the beginning of historical times. Additional and consistent information supporting this is that the fact that after decades of research in the region in several dozens of sites **[53]**, there is no evidence of corrals, guano deposits, llama caravanning camps (“nodes”), Andean artifacts such as pottery, art, religious figurines, or rocks of that origin, nor Andean tombs. Furthermore, there are no indications in the earliest historical chronicles of the region of llama caravanning or individuals of Andean origin, nor any linguistic evidence of any kind. It was also considered that the presence of dogs from wetland sites could be a product of exchange with the Andean region **[54]**, but today it has been proven that they were locally bred and extended continuously from the Paraná wetland to southern Brazil **[55]**. The only remaining evidence of some sort of Andean connection (which does not imply a direct connection) consists of extremely rare tiny metal fragments, recovered from a very small number of sites (**[55–58]**, and the references therein). In contrast, in over 40 sites that we have excavated over the past 15 years from these complex hunter-gatherer societies **[53]**, not a single piece of metal has been recovered. These excavations include several sites with extensive and randomly distributed sampling areas, ranging from 50 to 100 m², as well as numerous burial areas. There are also small beads made of green rocks that are more common and could have an Andean origin **[56,59, 60]**. However, these two records are based on small, portable artifacts typically associated with long trade networks, involving numerous episodes of direct person-to-person exchange over an extended period, with the gradual movement of goods **[56]**.

The results obtained in this study align fully with the known dispersal range of domesticated camelids and the extent of llama caravanning, as demonstrated by the extensive and detailed research in Andean archaeology. While some authors (e.g., **[4]**) have considered the possibility that certain historical sources suggest the presence of domesticated camelids in southwestern South America, these sources are more useful for generating hypotheses than for testing them. Moreover, many of these chronicles are geographically imprecise; in the 16^th^ century, the term ‘Río de la Plata’ referred to a much broader region, not limited to southeastern South America. The descriptions provided by travelers and sailors highlight the challenges they faced in identifying unfamiliar fauna, and it is clear that their taxonomic classifications do not correspond to modern standards **[61]**. While it is plausible to consider that mentions of “wild sheep” or “sheep of Peru” in geographically more precise and better-defined chronicles referring to southeastern South America may be referring to guanacos **[9]**, we have shown here that these wild camelids were indeed present in the northern Pampas plains and up to the Paraná Valley ecotone until the early historic period. In contrast, there is no archaeological evidence to support the presence of domesticated camelids in the region, nor such complex organizations as llama caravanning in southeastern South America.

## 5. Conclusions

By combining multiple lines of evidence from the archaeological record, stable isotope analysis, and paleogenomic data, we have confidently identified that all the analyzed camelids are guanacos. These populations were hunted throughout the Late Holocene until historical times. The particular record of guanacos in the Paraná valley sites began to form as a result of human activity around 700 BCE, and it is likely that this behavior began even earlier. No evidence of domesticated camelids has been identified in the region. This also allows us to currently rule out the presence of a complex, socially, politically, and economically impactful movement such as llama caravanning in southeastern South America. This study has also improved our understanding of the hunting strategies and capture areas associated with the guanacos recovered from the wetland ecotone sites, demonstrating the value of multi-analytical approaches in addressing interrelated questions and providing explanations based on a larger body of consistent data.

## 6. Acknowledgements and funding

Several grants funded this study: Ministerio de Cultura. Instituto Nacional de Antropología y Pensamiento Latinoamericano, Buenos Aires, Argentina. Consejo Nacional de Investigaciones Científicas y Técnicas, grants (PIPs) 11220110100565 / 11220150100482. MUR PRIN 2022 project 2022BC2Z5F ’Habits - Life-histories of prehistoric human groups in South America: a tale on environmental adaptations and subsistence economies. We would like to thank Eline Lorenzen for support of the palaeogenomic data generation and sequencing.

This study is part of the ASSA Program (www.assaprogram.com)

## Data availability

Raw sequencing reads for all ancient individuals are available under the following NCBI BioProject ID: PRJNA1160187

---

**Supplementary Figure 1.**
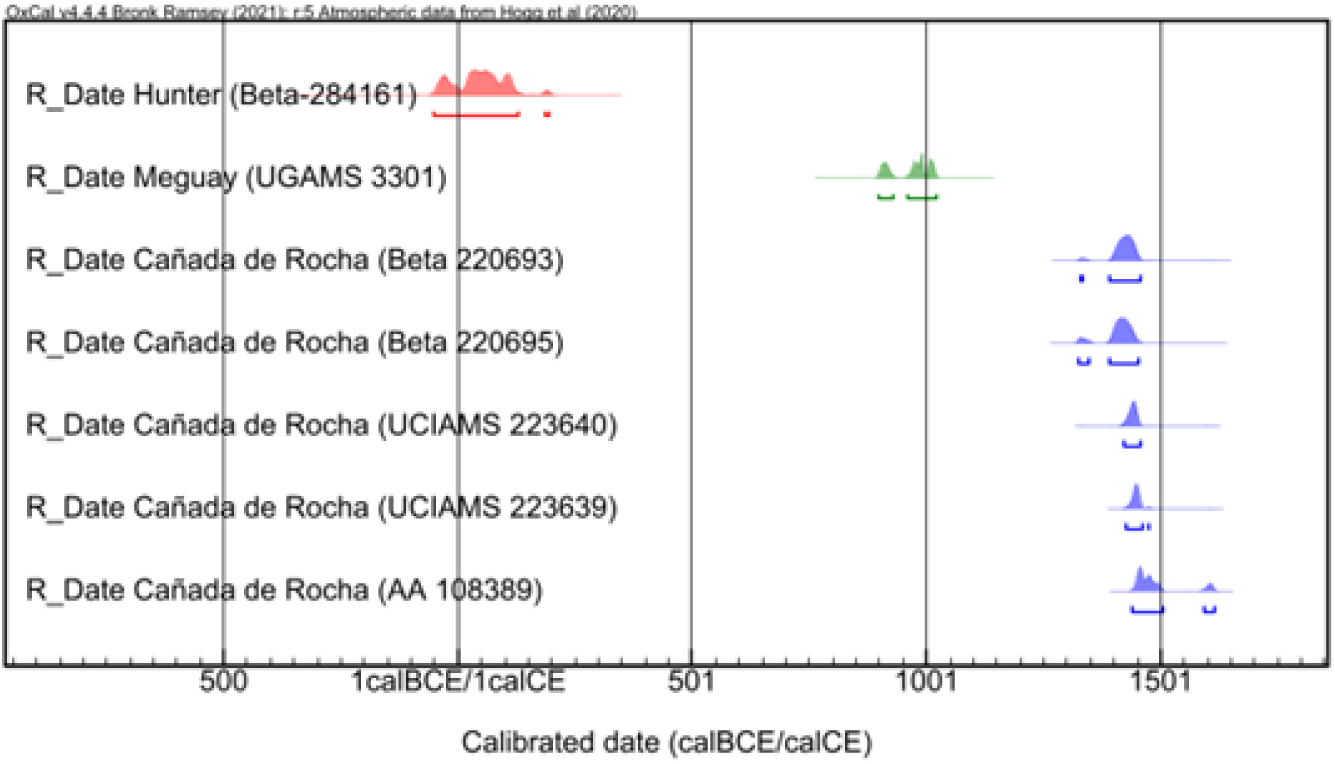
Calibrated date ranges of direct radiocarbon ages (AMS) on guanaco bones recovered from archaeological sites located at the Pampas plain.

**Supplementary Figure 2:**
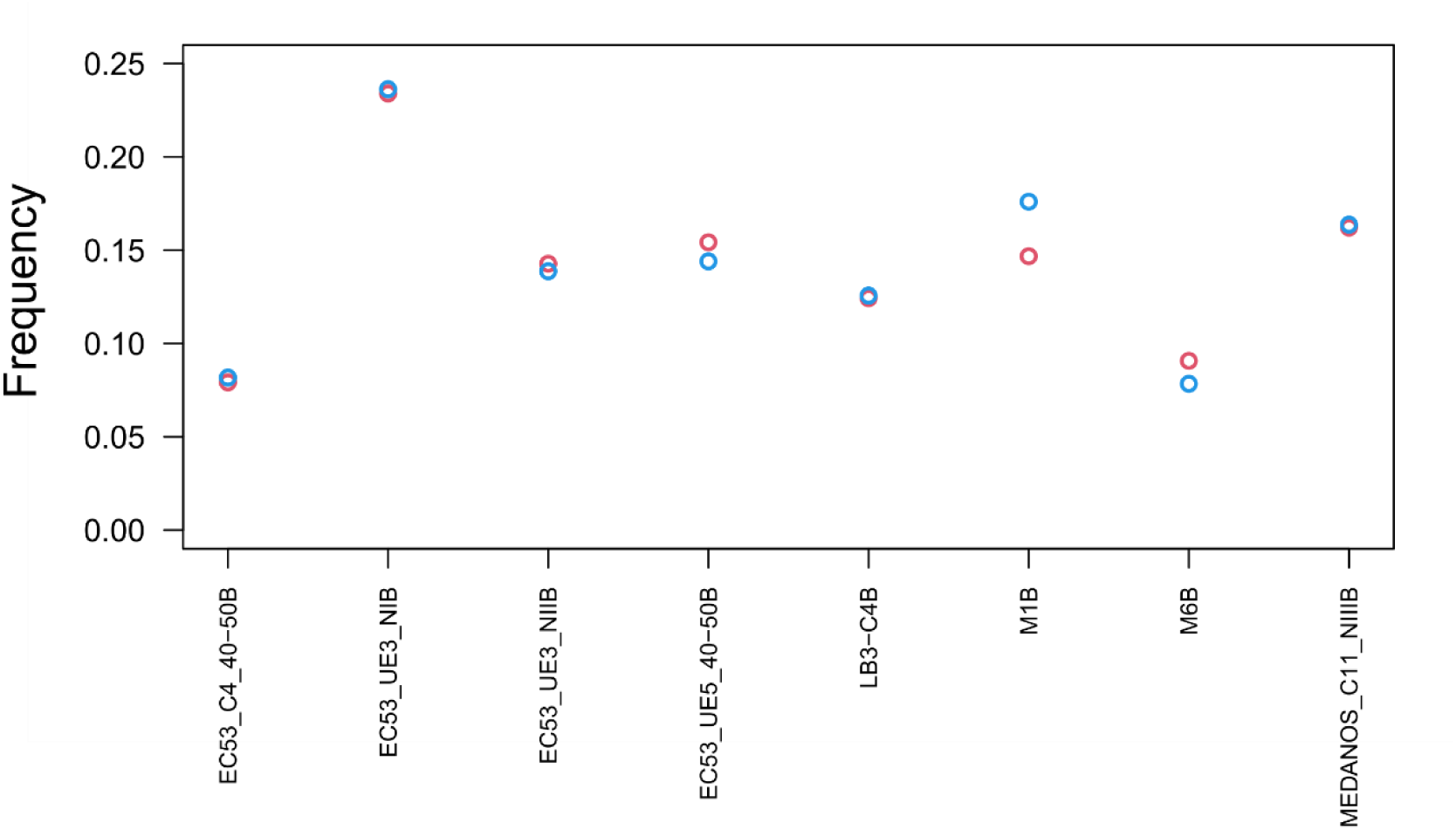
DNA damage results taken from Mapdamage for the eight individuals with sufficient data for sex determination. The frequency of C-T transitions on the first site from the 5-prime end of the read is shown in red, and G-A transitions on the first site from the 3-prime end of the read is shown in blue.

**Supplementary Figure 3:**
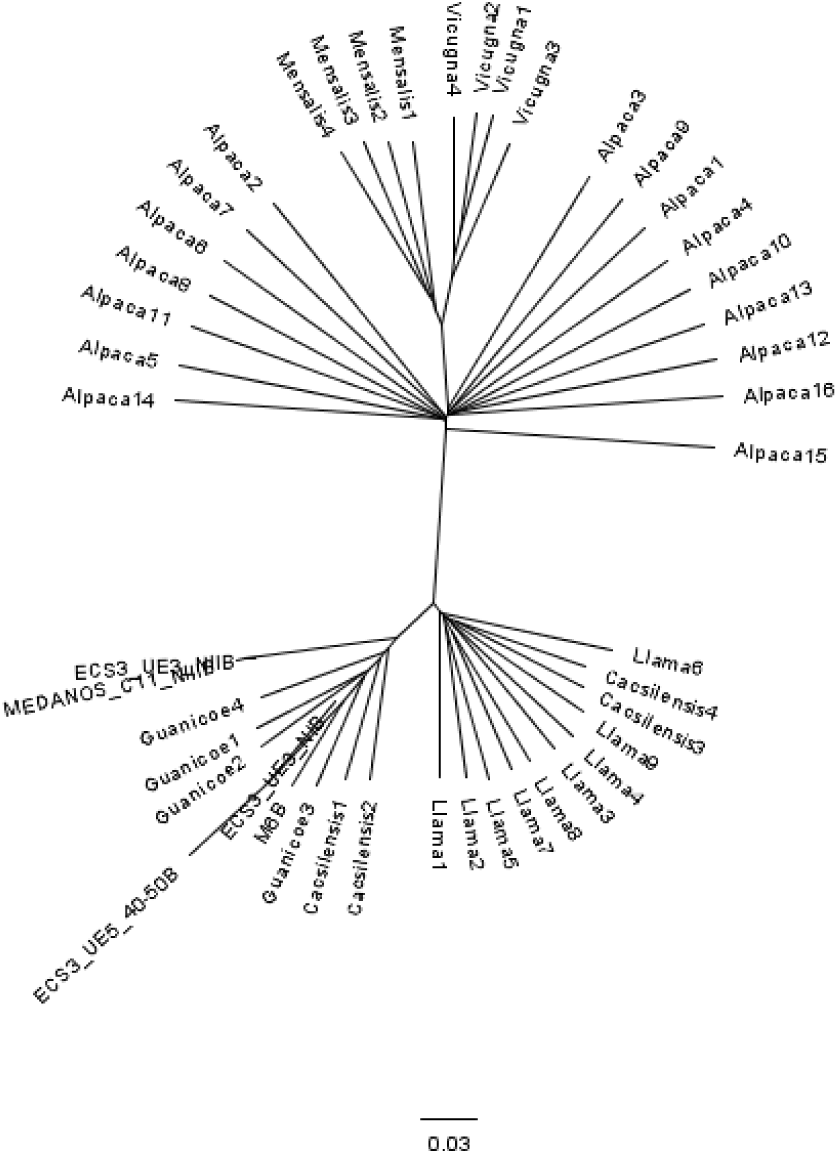
Neighbour joining tree constructed using the distance matrix computed in ANGSD using a pseudohaploid basecall. Branch lengths indicated identity by state distance.

**Supplementary Table 1.** Archaeological sites included in this study and located in the Pampas plain, 60-80 km from the ecotone with the wetland. All dated samples yielded a collagen content > 1%. In samples UCIAMS 223639 and 223640, the C/N ratio was measured, resulting in 3.23 and 3.24. The extended range from 1441 to 1615 CE of AMS AA 108389 is due to a plateau and a minor reversal in the calibration curve SHCal-20, The 84,4% range is 1440 to 1504 CE.

| Site | Location | Guanaco representation | Dated taxon | Lab. Code | Method | <sup>14</sup>C Age | s | Calibrated range CE (95%) | Reference |
| --- | --- | --- | --- | --- | --- | --- | --- | --- | --- |
| Hunter | Pampa plain | complete carcasses | *L. guanicoe* | Beta-284161 | AMS | 1990 | 40 | -51 to 196 | Loponte et al. (2010) |
| Meguay | Pampa plain | complete carcasses | *L. guanicoe* | UGAMS 3301 | AMS | 1120 | 20 | 897 -1022 | Loponte et al. (2010) |
| Cañada de Rocha | Pampa plain | complete carcasses | *L. guanicoe* | Beta 220693 | AMS | 540 | 40 | 1329-1458 | Toledo (2010) |
|  |  |  | *L. guanicoe* | Beta 220695 | AMS | 560 | 40 | 1324 - 1452 |  |
|  |  |  | *L. guanicoe* | UCIAMS 223639 | AMS | 485 | 20 | 1421 -1455 | Toledo (2021) |
|  |  |  | *L. guanicoe* | UCIAMS 223640 | AMS | 505 | 20 | 1426 - 1477 |  |
|  |  |  | *L. guanicoe* | AA 108389 | AMS | 452 | 24 | 1441 - 1615 | Buc & Loponte (2016) |

**Supplementary Table 2.**
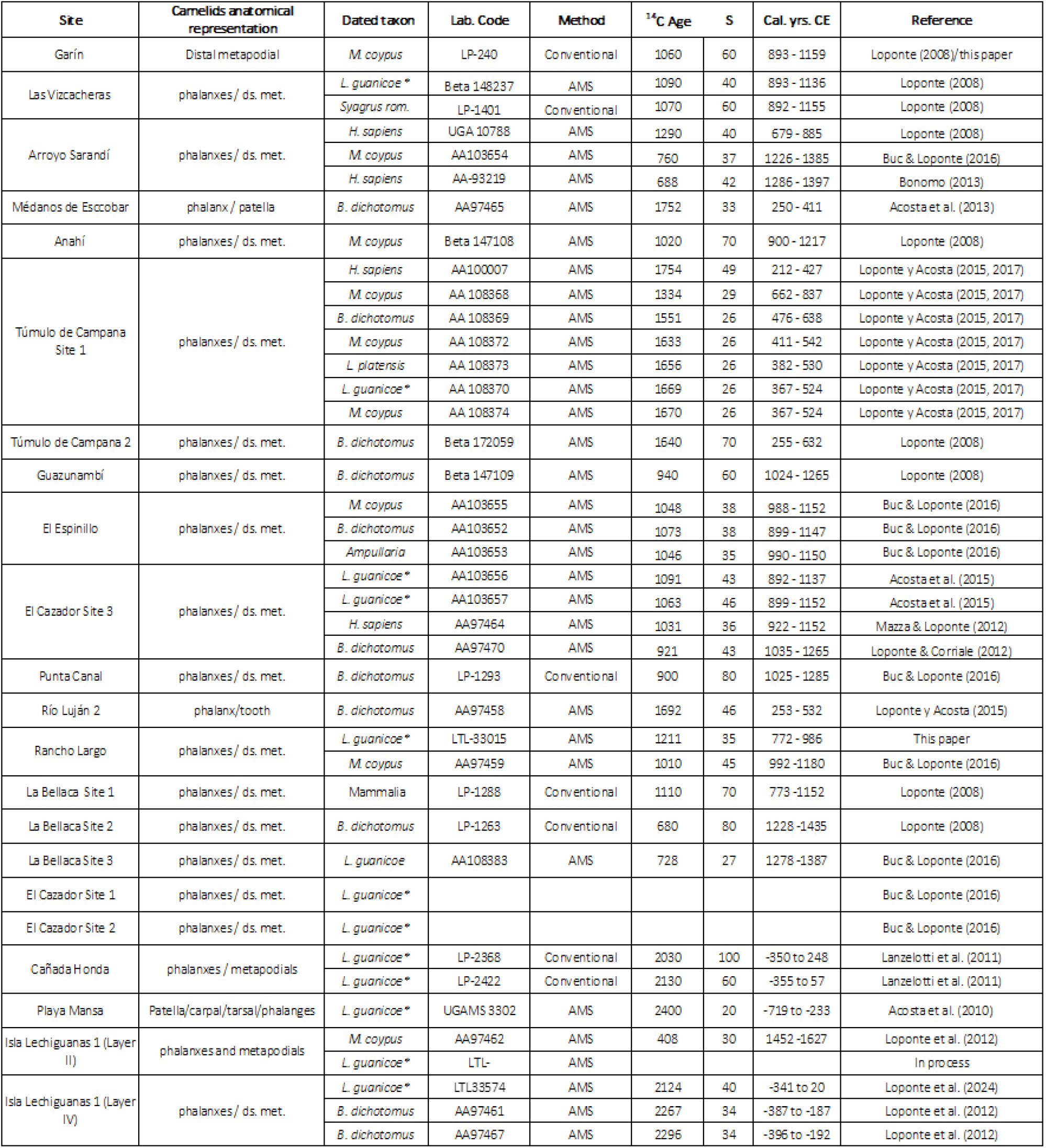
Archaeological sites included in this study located in the Lower Paraná wetland within the ecotone, or near the ecotone with the Pampas plain (Isla Lechiguanas 1). All dated samples yielded a collagen content > 1% and a carbon mass ≥ 1 mg, except at El Cazador site 1 and site 2. *Identification based on original authors (see references) and in the results of this study.

**Supplementary Table 3.** Fusion stages of metapodial condyles of camelids recovered from wetland ecotonal sites.

| Metapodial condyles (NISP) |  |  |
| --- | --- | --- |
| Sites | Fused | Unfused |
| Garín | 1 |  |
| Guazunambí | 3 | 1 |
| Túmulo Campana 2 |  | 1 |
| Rancho Largo | 12 | 4 |
| Sarandí | 3 | 1 |
| El Cazador 3 | 2 | 2 |
| El Cazador 2 | 1 |  |
| El Cazador 1 |  | 1 |
| El Espinillo | 2 | 2 |
| Punta Canal | 1 |  |
| La Bellaca 1 | 1 |  |
| La Bellaca 2 |  | 1 |
| La Bellaca 3 | 2 | 1 |
| Isla Lechiguanas1 (IV) | 1 |  |
| Total | 29 | 14 |
| % | 67.4 | 32.6 |

**Supplementary Table 4.**
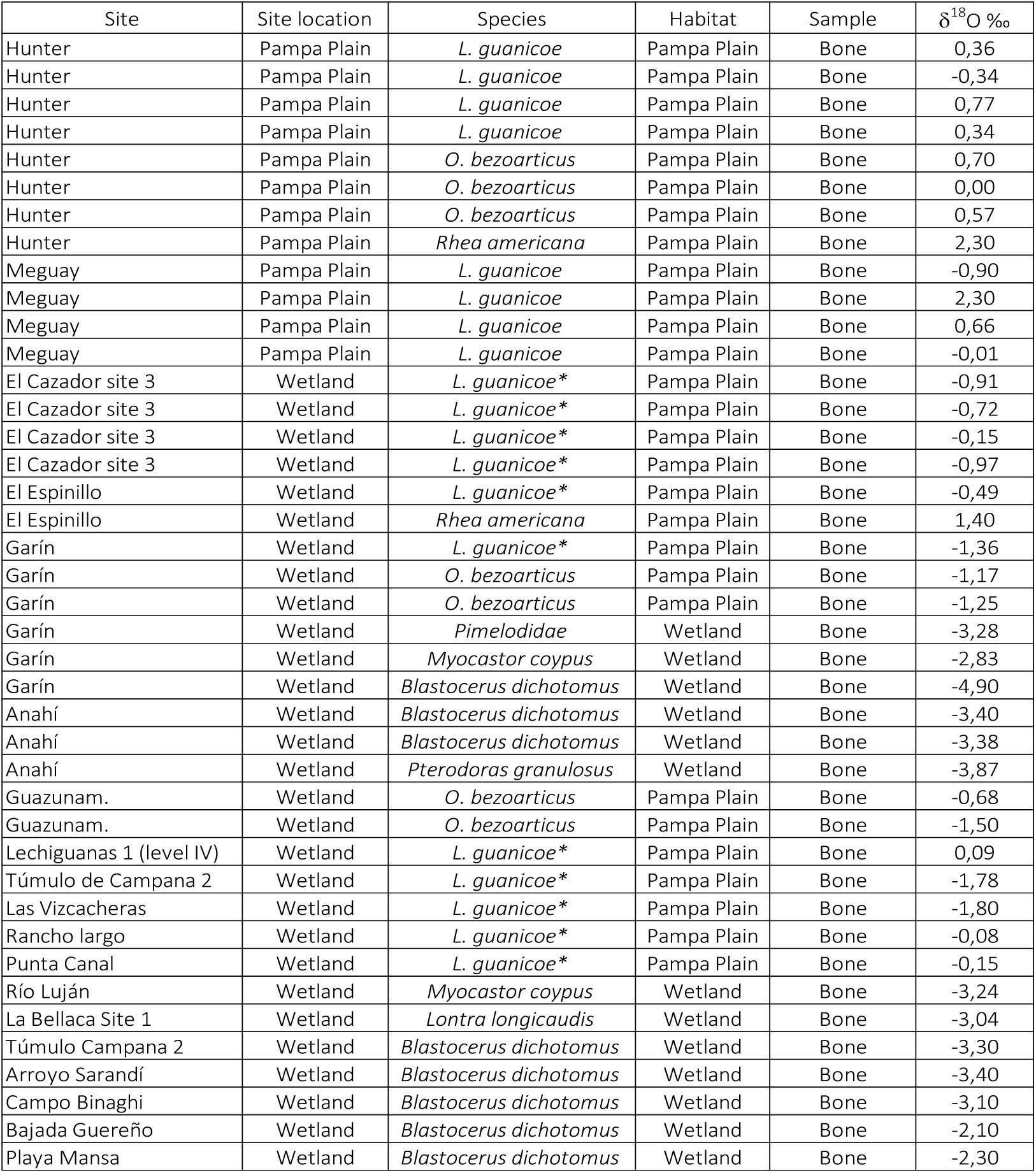
δ^18^O (‰) values (V-PDB) from the samples included in this study, originating from archaeological sites in the northern Pampas plain and the wetland ecotone adjacent to it along the Paraná River. * The results of this study show that these camelids correspond to *L. guanicoe* (see main text). Campo Binaghi site is located in the northern part of Santa Fe province, in the middle Paraná valley, but within the same δ^18^O ecozone as the lower Paraná valley.

**Supplementary Table 5.** Values of δ^13^C (‰) (V-PDB) from camelids and deer from archaeological sites in the northern Pampean region and the lower Paraná wetland ecotone. *Determination based on the results obtained in this study.

| Site | Site location | Species | Habitat | Sample | $\delta^{13}\text{C}$ (‰) |
| --- | --- | --- | --- | --- | --- |
| Hunter | Pampa plain | *L. guanicoe* | Pampa plain | Bone | -8.7 |
| Hunter | Pampa plain | *L. guanicoe* | Pampa plain | Bone | -8.5 |
| Hunter | Pampa plain | *L. guanicoe* | Pampa plain | Bone | -10.2 |
| Hunter | Pampa plain | *L. guanicoe* | Pampa plain | Bone | -7.7 |
| Hunter | Pampa plain | *O. bezoarticus* | Pampa plain | Bone | 11.4 |
| Hunter | Pampa plain | *O. bezoarticus* | Pampa plain | Bone | 10.0 |
| Hunter | Pampa plain | *O. bezoarticus* | Pampa plain | Bone | 10.5 |
| Meguay | Pampa plain | *L. guanicoe* | Pampa plain | Bone | -9.2 |
| Meguay | Pampa plain | *L. guanicoe* | Pampa plain | Bone | -9.0 |
| Meguay | Pampa plain | *L. guanicoe* | Pampa plain | Bone | -10.3 |
| Meguay | Pampa plain | *L. guanicoe* | Pampa plain | Bone | -8.6 |
| El Cazador sitio 3 | Wetland | *L. guanicoe* * | Pampa plain | Bone | -8.8 |
| El Cazador sitio 3 | Wetland | *L. guanicoe* * | Pampa plain | Bone | -9.1 |
| El Cazador sitio 3 | Wetland | *L. guanicoe* * | Pampa plain | Bone | -9.4 |
| El Cazador sitio 3 | Wetland | *L. guanicoe* * | Pampa plain | Bone | -10.9 |
| El Cazador sitio 3 | Wetland | *L. guanicoe* * | Pampa plain | Bone | -9.0 |
| El Espinillo | Wetland | *L. guanicoe* * | Pampa plain | Bone | -8.3 |
| Garín | Wetland | *L. guanicoe* * | Pampa plain | Bone | -13.1 |
| Lechiguanas 1 (level IV) | Wetland | *L. guanicoe* * | Pampa plain | Bone | -9.2 |
| Túmulo de Campana 2 | Wetland | *L. guanicoe* * | Pampa plain | Bone | -10.2 |
| Vizcach. | Wetland | *L. guanicoe* * | Pampa plain | Bone | -9.5 |
| Rancho largo | Wetland | *L. guanicoe* * | Pampa plain | Bone | -9.7 |
| Punta Canal | Wetland | *L. guanicoe* * | Pampa plain | Bone | -9.5 |

**Supplementary Table 6.**
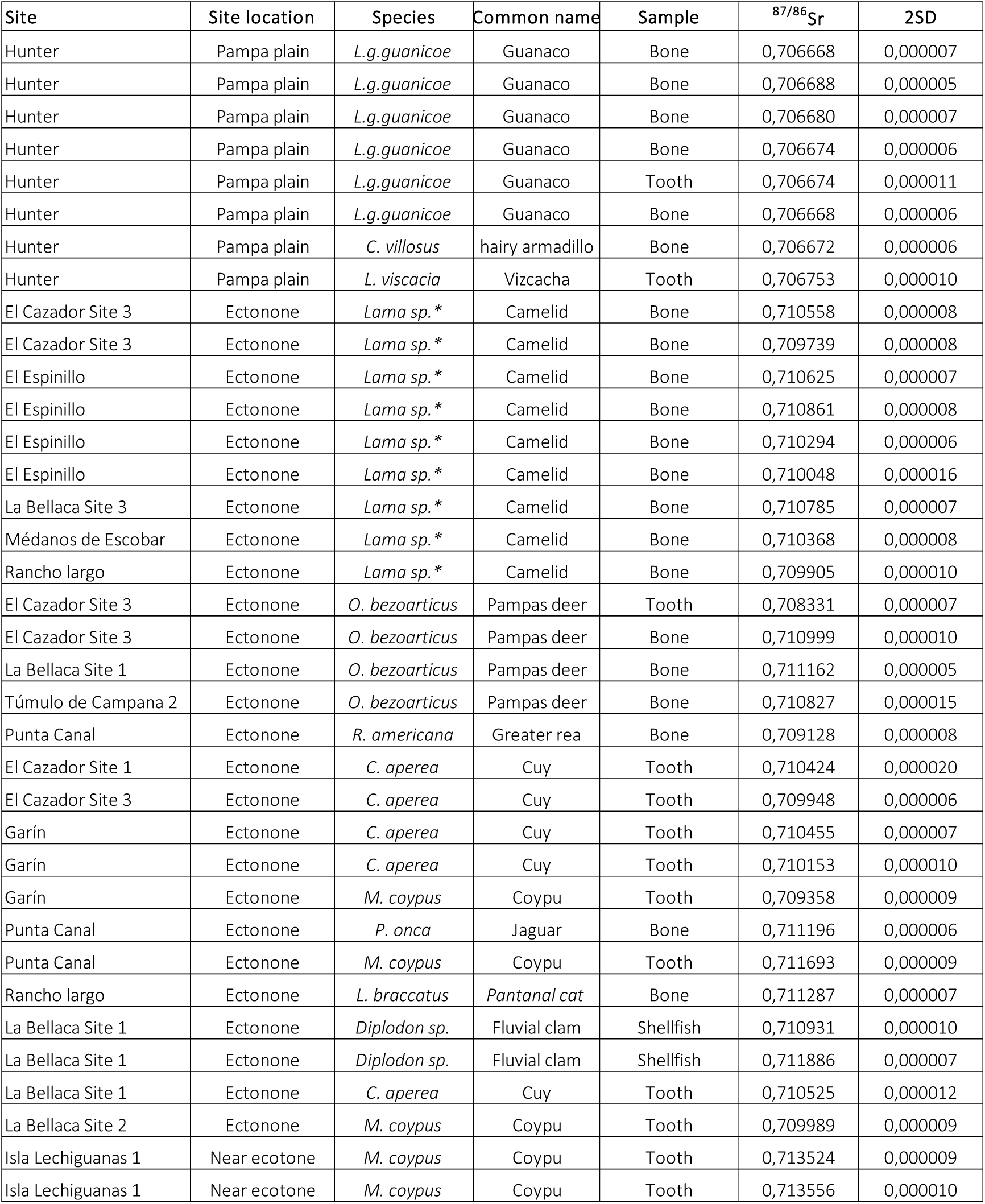
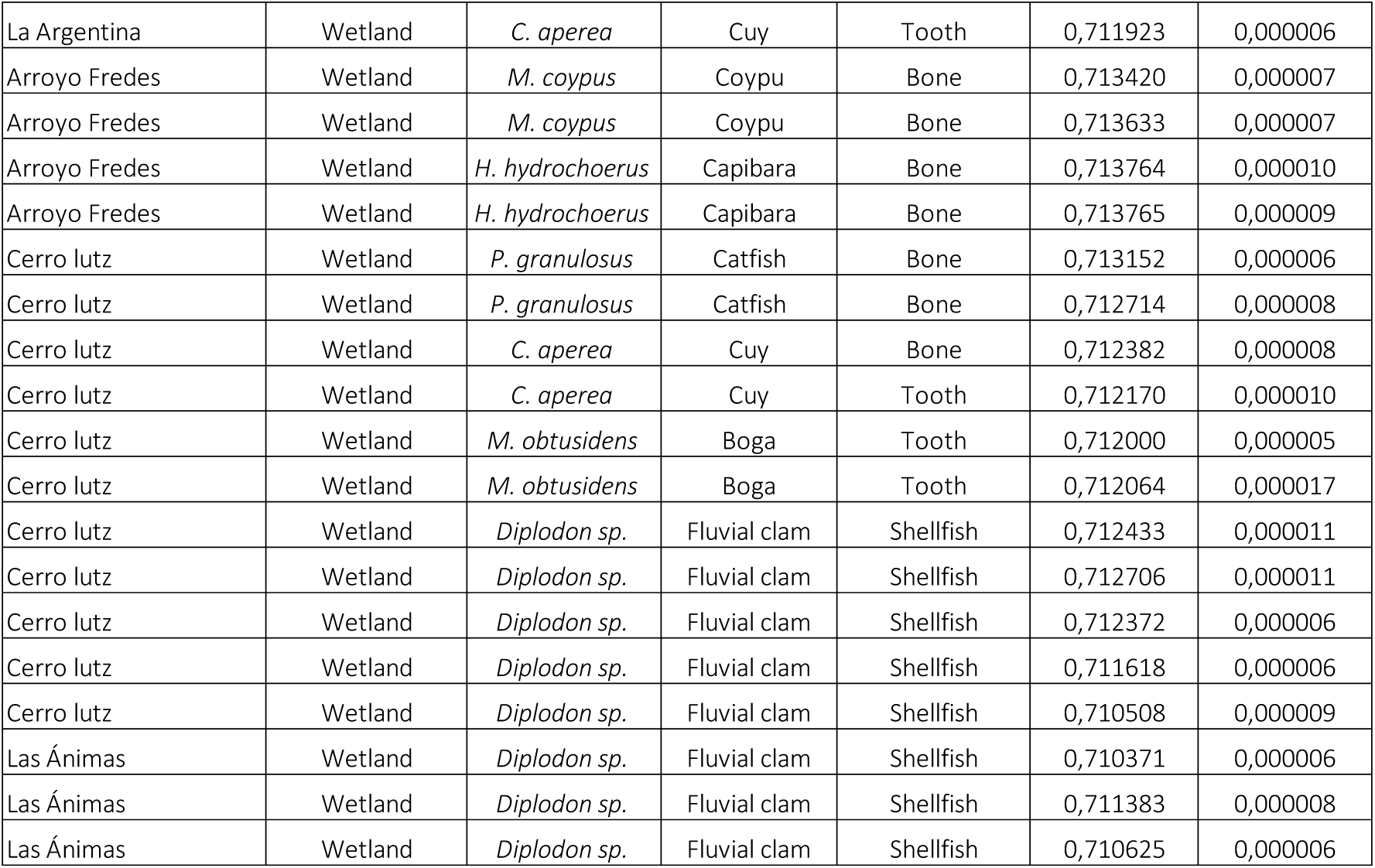
Values of ^87^Sr/^86^Sr from taxa recovered at northern Pampa plains and lower Paraná wetland. * Identified as *L. guanicoe* after this study.

**Supplementary Table 7.**
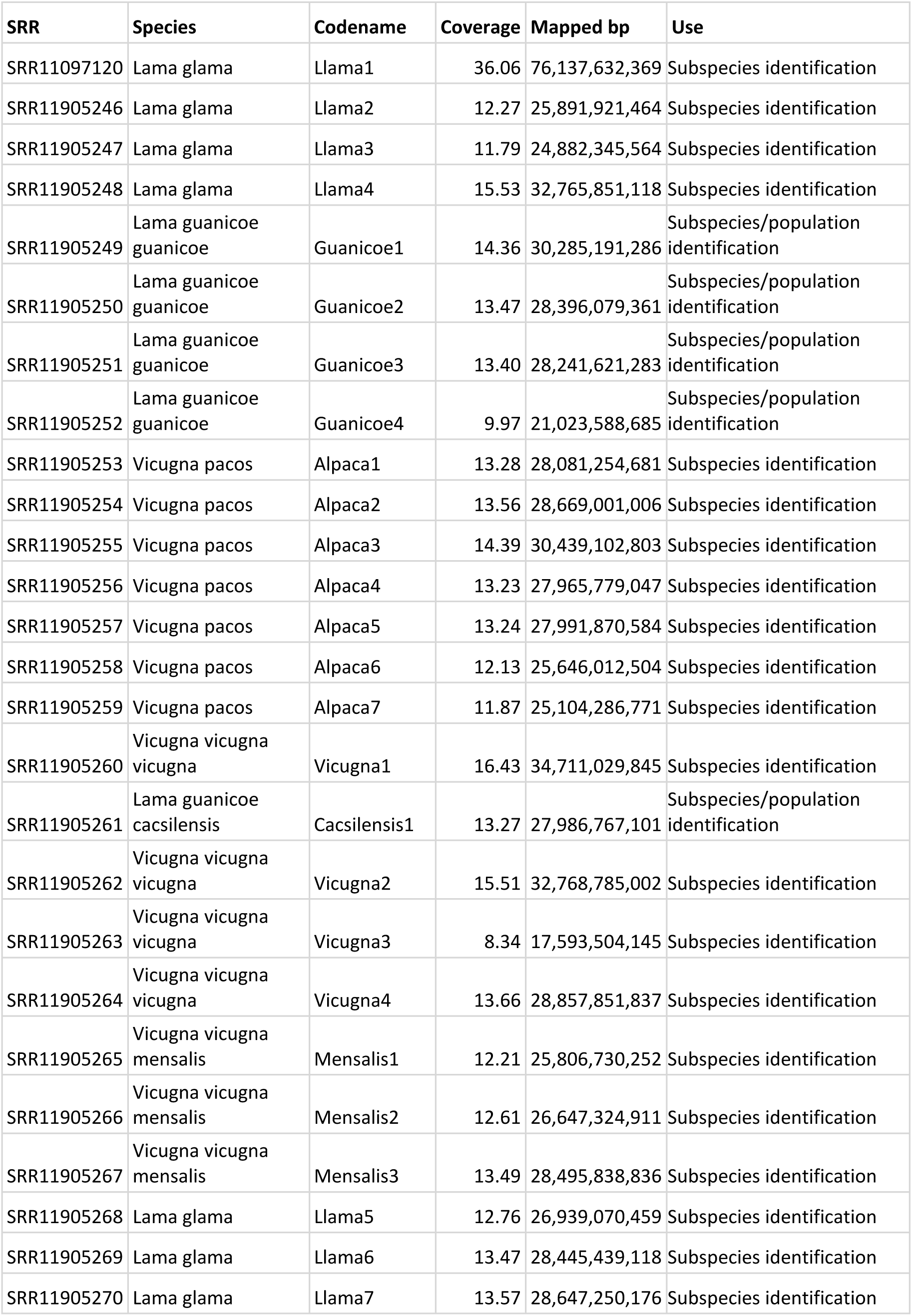
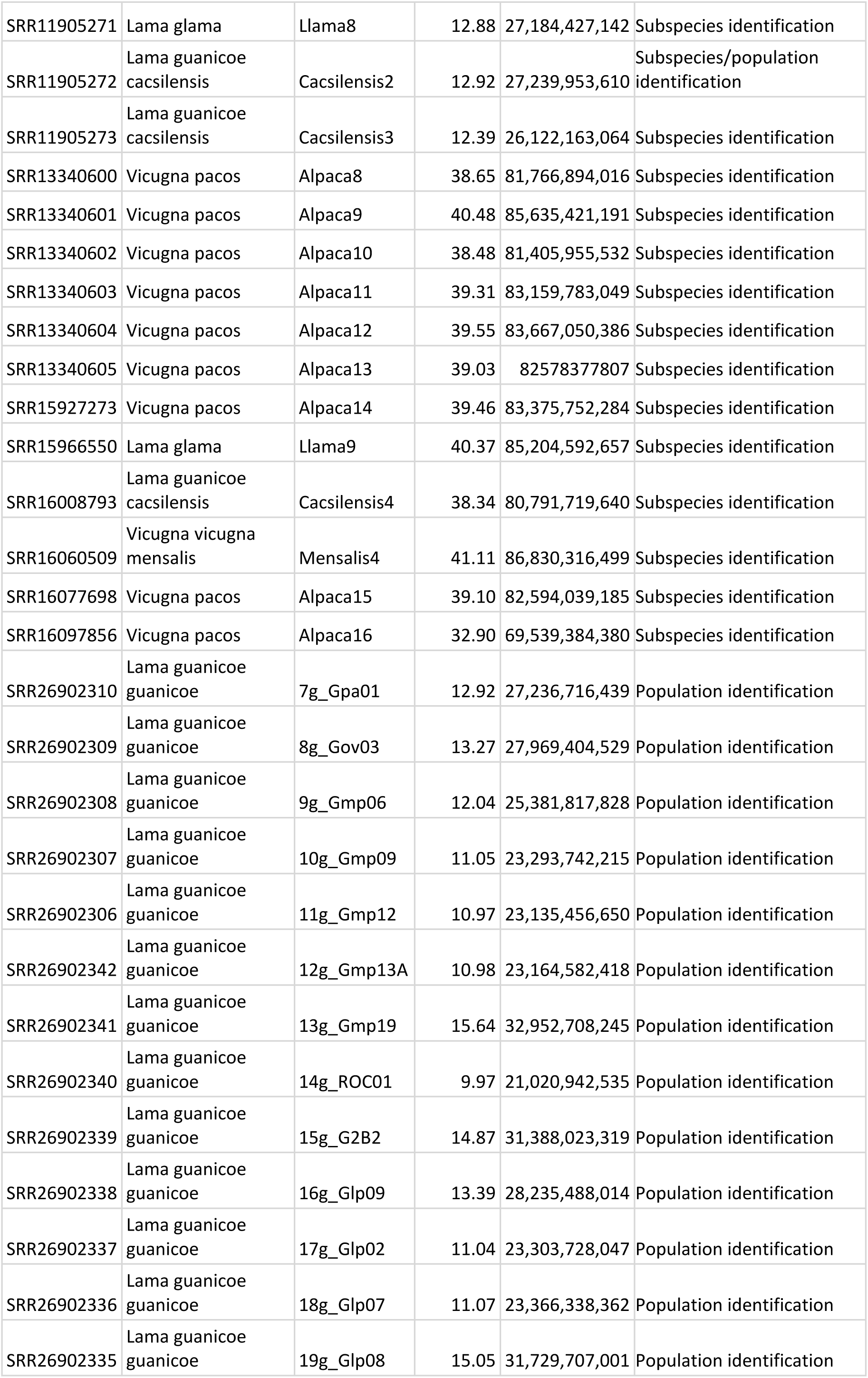
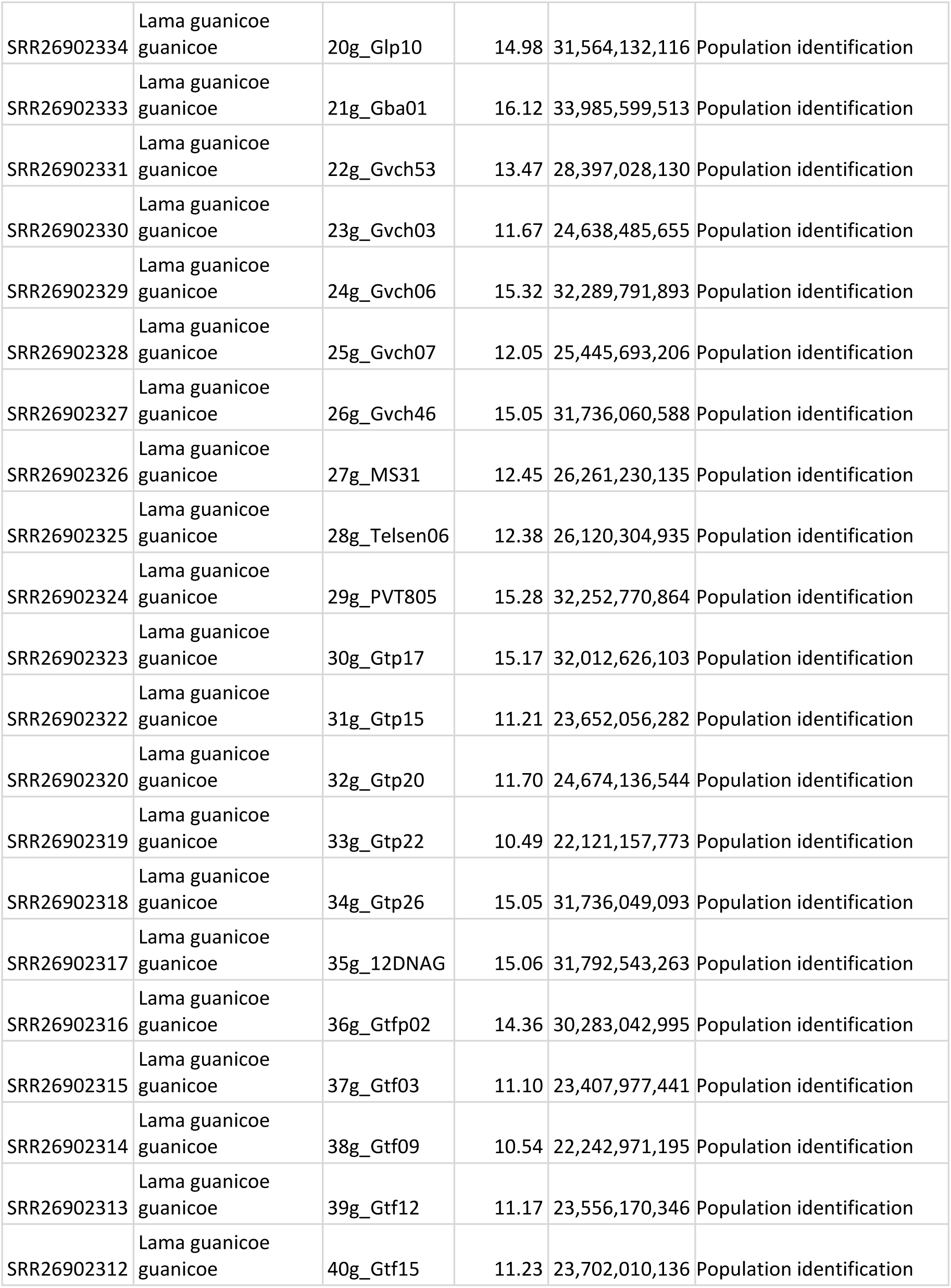
Accession codes, sample information and mapping statistics for the modern genomic data

| SRR | Species | Codename | Coverage | Mapped bp | Use |
| --- | --- | --- | --- | --- | --- |
| SRR11097120 | Lama glama | Llama1 | 36.06 | 76,137,632,369 | Subspecies identification |
| SRR11905246 | Lama glama | Llama2 | 12.27 | 25,891,921,464 | Subspecies identification |
| SRR11905247 | Lama glama | Llama3 | 11.79 | 24,882,345,564 | Subspecies identification |
| SRR11905248 | Lama glama | Llama4 | 15.53 | 32,765,851,118 | Subspecies identification |
| SRR11905249 | Lama guanicoe<br>guanicoe | Guanicoe1 | 14.36 | 30,285,191,286 | Subspecies/population<br>identification |
| SRR11905250 | Lama guanicoe<br>guanicoe | Guanicoe2 | 13.47 | 28,396,079,361 | Subspecies/population<br>identification |
| SRR11905251 | Lama guanicoe<br>guanicoe | Guanicoe3 | 13.40 | 28,241,621,283 | Subspecies/population<br>identification |
| SRR11905252 | Lama guanicoe<br>guanicoe | Guanicoe4 | 9.97 | 21,023,588,685 | Subspecies/population<br>identification |
| SRR11905253 | Vicugna pacos | Alpaca1 | 13.28 | 28,081,254,681 | Subspecies identification |
| SRR11905254 | Vicugna pacos | Alpaca2 | 13.56 | 28,669,001,006 | Subspecies identification |
| SRR11905255 | Vicugna pacos | Alpaca3 | 14.39 | 30,439,102,803 | Subspecies identification |
| SRR11905256 | Vicugna pacos | Alpaca4 | 13.23 | 27,965,779,047 | Subspecies identification |
| SRR11905257 | Vicugna pacos | Alpaca5 | 13.24 | 27,991,870,584 | Subspecies identification |
| SRR11905258 | Vicugna pacos | Alpaca6 | 12.13 | 25,646,012,504 | Subspecies identification |
| SRR11905259 | Vicugna pacos | Alpaca7 | 11.87 | 25,104,286,771 | Subspecies identification |
| SRR11905260 | Vicugna vicugna<br>vicugna | Vicugna1 | 16.43 | 34,711,029,845 | Subspecies identification |
| SRR11905261 | Lama guanicoe<br>cacsilensis | Cacsilensis1 | 13.27 | 27,986,767,101 | Subspecies/population<br>identification |
| SRR11905262 | Vicugna vicugna<br>vicugna | Vicugna2 | 15.51 | 32,768,785,002 | Subspecies identification |
| SRR11905263 | Vicugna vicugna<br>vicugna | Vicugna3 | 8.34 | 17,593,504,145 | Subspecies identification |
| SRR11905264 | Vicugna vicugna<br>vicugna | Vicugna4 | 13.66 | 28,857,851,837 | Subspecies identification |
| SRR11905265 | Vicugna vicugna<br>mensalis | Mensalis1 | 12.21 | 25,806,730,252 | Subspecies identification |
| SRR11905266 | Vicugna vicugna<br>mensalis | Mensalis2 | 12.61 | 26,647,324,911 | Subspecies identification |
| SRR11905267 | Vicugna vicugna<br>mensalis | Mensalis3 | 13.49 | 28,495,838,836 | Subspecies identification |
| SRR11905268 | Lama glama | Llama5 | 12.76 | 26,939,070,459 | Subspecies identification |
| SRR11905269 | Lama glama | Llama6 | 13.47 | 28,445,439,118 | Subspecies identification |
| SRR11905270 | Lama glama | Llama7 | 13.57 | 28,647,250,176 | Subspecies identification |
| SRR11905271 | Lama glama | Llama8 | 12.88 | 27,184,427,142 | Subspecies identification |
| SRR11905272 | Lama guanicoe cacsilensis | Cacsilensis2 | 12.92 | 27,239,953,610 | Subspecies/population identification |
| SRR11905273 | Lama guanicoe cacsilensis | Cacsilensis3 | 12.39 | 26,122,163,064 | Subspecies identification |
| SRR13340600 | Vicugna pacos | Alpaca8 | 38.65 | 81,766,894,016 | Subspecies identification |
| SRR13340601 | Vicugna pacos | Alpaca9 | 40.48 | 85,635,421,191 | Subspecies identification |
| SRR13340602 | Vicugna pacos | Alpaca10 | 38.48 | 81,405,955,532 | Subspecies identification |
| SRR13340603 | Vicugna pacos | Alpaca11 | 39.31 | 83,159,783,049 | Subspecies identification |
| SRR13340604 | Vicugna pacos | Alpaca12 | 39.55 | 83,667,050,386 | Subspecies identification |
| SRR13340605 | Vicugna pacos | Alpaca13 | 39.03 | 82578377807 | Subspecies identification |
| SRR15927273 | Vicugna pacos | Alpaca14 | 39.46 | 83,375,752,284 | Subspecies identification |
| SRR15966550 | Lama glama | Llama9 | 40.37 | 85,204,592,657 | Subspecies identification |
| SRR16008793 | Lama guanicoe cacsilensis | Cacsilensis4 | 38.34 | 80,791,719,640 | Subspecies identification |
| SRR16060509 | Vicugna vicugna mensalis | Mensalis4 | 41.11 | 86,830,316,499 | Subspecies identification |
| SRR16077698 | Vicugna pacos | Alpaca15 | 39.10 | 82,594,039,185 | Subspecies identification |
| SRR16097856 | Vicugna pacos | Alpaca16 | 32.90 | 69,539,384,380 | Subspecies identification |
| SRR26902310 | Lama guanicoe guanicoe | 7g_Gpa01 | 12.92 | 27,236,716,439 | Population identification |
| SRR26902309 | Lama guanicoe guanicoe | 8g_Gov03 | 13.27 | 27,969,404,529 | Population identification |
| SRR26902308 | Lama guanicoe guanicoe | 9g_Gmp06 | 12.04 | 25,381,817,828 | Population identification |
| SRR26902307 | Lama guanicoe guanicoe | 10g_Gmp09 | 11.05 | 23,293,742,215 | Population identification |
| SRR26902306 | Lama guanicoe guanicoe | 11g_Gmp12 | 10.97 | 23,135,456,650 | Population identification |
| SRR26902342 | Lama guanicoe guanicoe | 12g_Gmp13A | 10.98 | 23,164,582,418 | Population identification |
| SRR26902341 | Lama guanicoe guanicoe | 13g_Gmp19 | 15.64 | 32,952,708,245 | Population identification |
| SRR26902340 | Lama guanicoe guanicoe | 14g_ROC01 | 9.97 | 21,020,942,535 | Population identification |
| SRR26902339 | Lama guanicoe guanicoe | 15g_G2B2 | 14.87 | 31,388,023,319 | Population identification |
| SRR26902338 | Lama guanicoe guanicoe | 16g_Glp09 | 13.39 | 28,235,488,014 | Population identification |
| SRR26902337 | Lama guanicoe guanicoe | 17g_Glp02 | 11.04 | 23,303,728,047 | Population identification |
| SRR26902336 | Lama guanicoe guanicoe | 18g_Glp07 | 11.07 | 23,366,338,362 | Population identification |
| SRR26902335 | Lama guanicoe guanicoe | 19g_Glp08 | 15.05 | 31,729,707,001 | Population identification |
| SRR26902334 | Lama guanicoe<br>guanicoe | 20g_Glp10 | 14.98 | 31,564,132,116 | Population identification |
| SRR26902333 | Lama guanicoe<br>guanicoe | 21g_Gba01 | 16.12 | 33,985,599,513 | Population identification |
| SRR26902331 | Lama guanicoe<br>guanicoe | 22g_Gvch53 | 13.47 | 28,397,028,130 | Population identification |
| SRR26902330 | Lama guanicoe<br>guanicoe | 23g_Gvch03 | 11.67 | 24,638,485,655 | Population identification |
| SRR26902329 | Lama guanicoe<br>guanicoe | 24g_Gvch06 | 15.32 | 32,289,791,893 | Population identification |
| SRR26902328 | Lama guanicoe<br>guanicoe | 25g_Gvch07 | 12.05 | 25,445,693,206 | Population identification |
| SRR26902327 | Lama guanicoe<br>guanicoe | 26g_Gvch46 | 15.05 | 31,736,060,588 | Population identification |
| SRR26902326 | Lama guanicoe<br>guanicoe | 27g_MS31 | 12.45 | 26,261,230,135 | Population identification |
| SRR26902325 | Lama guanicoe<br>guanicoe | 28g_Telsen06 | 12.38 | 26,120,304,935 | Population identification |
| SRR26902324 | Lama guanicoe<br>guanicoe | 29g_PVT805 | 15.28 | 32,252,770,864 | Population identification |
| SRR26902323 | Lama guanicoe<br>guanicoe | 30g_Gtp17 | 15.17 | 32,012,626,103 | Population identification |
| SRR26902322 | Lama guanicoe<br>guanicoe | 31g_Gtp15 | 11.21 | 23,652,056,282 | Population identification |
| SRR26902320 | Lama guanicoe<br>guanicoe | 32g_Gtp20 | 11.70 | 24,674,136,544 | Population identification |
| SRR26902319 | Lama guanicoe<br>guanicoe | 33g_Gtp22 | 10.49 | 22,121,157,773 | Population identification |
| SRR26902318 | Lama guanicoe<br>guanicoe | 34g_Gtp26 | 15.05 | 31,736,049,093 | Population identification |
| SRR26902317 | Lama guanicoe<br>guanicoe | 35g_12DNAG | 15.06 | 31,792,543,263 | Population identification |
| SRR26902316 | Lama guanicoe<br>guanicoe | 36g_Gtfp02 | 14.36 | 30,283,042,995 | Population identification |
| SRR26902315 | Lama guanicoe<br>guanicoe | 37g_Gtf03 | 11.10 | 23,407,977,441 | Population identification |
| SRR26902314 | Lama guanicoe<br>guanicoe | 38g_Gtf09 | 10.54 | 22,242,971,195 | Population identification |
| SRR26902313 | Lama guanicoe<br>guanicoe | 39g_Gtf12 | 11.17 | 23,556,170,346 | Population identification |
| SRR26902312 | Lama guanicoe<br>guanicoe | 40g_Gtf15 | 11.23 | 23,702,010,136 | Population identification |

**Supplementary Table 8.** Sample information, mapping statistics and sex identification for the palaeogenomic data

| Sample ID | Tissue | Type sample | Archaeological site | Identified species | Recovered from | Raw reads | Trimmed reads | Mapped unique reads | Coverage | Mapped bp | Genetic sex | SeXY mean | Average read length |
| --- | --- | --- | --- | --- | --- | --- | --- | --- | --- | --- | --- | --- | --- |
| ECS3_C4_40-50 | Bone (metapodial) | Bone instrument | El Cazador Site 3 | ? | Ecotone (wetland) | 14174659 | 8465314 | 12470 | 0,0003 | 676.835 | F | 1,05 | 54 |
| ECS3_UE3_NI | Bone (metapodial) | Bone instrument | El Cazador Site 3 | ? | Ecotone (wetland) | 7936570 | 5860395 | 27242 | 0,0008 | 1.574.375 | F | 0,91 | 57 |
| ECS3_UE3_NII | Bone (metapodial) | Bone instrument | El Cazador Site 3 | ? | Ecotone (wetland) | 13872743 | 9597944 | 32501 | 0,0009 | 1.794.536 | F | 0,94 | 55 |
| ECS3_UE5_40-50 | Bone (metapodial) | Bone instrument | El Cazador Site 3 | ? | Ecotone (wetland) | 5233901 | 3582002 | 79512 | 0,0017 | 3.488.949 | F | 1,13 | 44 |
| ESP_C17 | Bone (phalanx) | Food discard | El Espinillo | ? | Ecotone (wetland) | 11761762 | 6009685 | 3227 | 0,0001 | 125.497 | NA | NA | 38 |
| ESP_C14 | Bone (metapodial) | Bone instrument | El Espinillo | ? | Ecotone (wetland) | 5456800 | 3454041 | 2066 | 0,0001 | 123.303 | NA | NA | 59 |
| LB3-C4 | Bone (phalanx) | Food discard | La Bellaca site 3 | ? | Ecotone (wetland) | 15571757 | 9431542 | 17321 | 0,0004 | 813.027 | M | 0,50 | 46 |
| MEDANOS_C11_NIII | Bone (patella) | Food discard | Médanos Escobar | ? | Ecotone (wetland) | 13187792 | 8732863 | 22271 | 0,0006 | 1.232.363 | F | 1,02 | 55 |
| H1 | Bone (phalanx) | Food discard | Hunter | *L. guanicoe* | Pampas plain | 5310674 | 3901229 | 247 | 0,0000 | 11.625 | NA | NA | 47 |
| H3 | Bone (phalanx) | Food discard | Hunter | *L. guanicoe* | Pampas plain | 4253337 | 3394019 | 118 | 0,0000 | 5.660 | NA | NA | 48 |
| H4 | Bone (metapodial) | Food discard | Hunter | *L. guanicoe* | Pampas plain | 3374363 | 2547299 | 167 | 0,0000 | 8.124 | NA | NA | 48 |
| H5 | Bone (metapodial) | Food discard | Hunter | *L. guanicoe* | Pampas plain | 7044792 | 5327501 | 138 | 0,0000 | 7.845 | NA | NA | 56 |
| H6 | Bone (phalanx) | Food discard | Hunter | *L. guanicoe* | Pampas plain | 10851028 | 8260228 | 524 | 0,0000 | 25.444 | NA | NA | 48 |
| H7 | Bone (metapodial) | Food discard | Hunter | *L. guanicoe* | Pampas plain | 1189748 | 811232 | 47 | 0,0000 | 2.509 | NA | NA | 53 |
| H8 | Bone (metapodial) | Food discard | Hunter | *L. guanicoe* | Pampas plain | 4177909 | 3268788 | 246 | 0,0000 | 13.930 | NA | NA | 56 |
| H9 | molar) | Food discard | Hunter | *L. guanicoe* | Pampas plain | 6634514 | 3753068 | 882 | 0,0000 | 36.019 | NA | NA | 40 |
| M1 | Bone (phalanx) | Food discard | Meguay | *L. guanicoe* | Pampas plain | 7536733 | 5075900 | 6472 | 0,0001 | 286.576 | M | 0,45 | 44 |
| M2 | Bone (phalanx) | Food discard | Meguay | *L. guanicoe* | Pampas plain | 14795388 | 10419113 | 453 | 0,0000 | 18.702 | NA | NA | 41 |
| M3 | Bone (phalanx) | Food discard | Meguay | *L. guanicoe* | Pampas plain | 13174874 | 13174874 | 481 | 0,0000 | 21.928 | NA | NA | 46 |
| M4 | Bone (metapodial) | Food discard | Meguay | *L. guanicoe* | Pampas plain | 8664678 | 5413891 | 808 | 0,0000 | 36.340 | NA | NA | 45 |
| M5 | Bone (metapodial) | Food discard | Meguay | *L. guanicoe* | Pampas plain | 5575669 | 4224249 | 202 | 0,0000 | 8.894 | NA | NA | 44 |
| M6 | Bone (metapodial) | Food discard | Meguay | *L. guanicoe* | Pampas plain | 13030109 | 9449971 | 30648 | 0,0007 | 1.515.481 | F | 1,03 | 49 |

## Supplementary Text

### 1. Archaeological context

The Paraná River wetland is an exceptionally rich environment in terms of fauna and biodiversity, supporting numerous gregarious species closely associated with water bodies and subtropical humid environments. Among these, freshwater fish were the most intensively exploited faunal resource by local indigenous population, followed by *Myocastor coypus*, *Cavia aperea*, and *Diplodon* sp. (Table 1). The wetland is also home to the marsh deer (*Blastocerus dichotomus*), a mammal weighing up to 150 kg, whose distribution is closely linked to this swampy environment, as it did not inhabit the adjacent plains (Loponte & Corriale, 2019). This species was heavily exploited by local hunters throughout the Late Holocene. There is also a small quantity of Pampa deer bones (*Ozotoceros bezoarticus*), as although this small deer (⁓ 25 kg) is typical of the adjacent Pampean plain, its ethological and ecological plasticity allowed it to occasionally venture into the flooded plains of the wetland, where it was eventually hunted (Loponte & Corriale, 2019). In contrast, bones of *Rhea americana* (great er rhea), a species typical of the adjacent plains, are extremely scarce, as is the case with camelid remains (Table 1). The identification of both rhea and camelid bones, like all rare taxa in archaeofaunal assemblages, is largely dependent on sample size (Lyman, 1995). At wetland ecotone sites, these species are generally only detected in collections with more than 5,000 bone fragments. Complete carcasses of wetland species, including marsh deer, were transported intact to residential sites for processing. Although the pampas deer is not native to wetlands, it was a frequent visitor to the floodplain grasslands in this environment, accessing them directly from the Pampas plain. Its anatomical representation at ecotone sites indicates that it was brought as whole prey to the residential camps. Bone preservation at these sites is excellent, with no evidence of preservation bias related to bone mineral density. Hyoid bones, intact proximal humeral epiphyses from mammals, flat and thin fish bones with low mineral density, such as opercular bones, and even delicate fish scales were frequently recovered. The weathering stages of the bone assemblages range from 0 to 3 on the Behrensmeyer scale (1978), with most bones falling into stage 1 (Loponte, 2008).

**Table 1.**
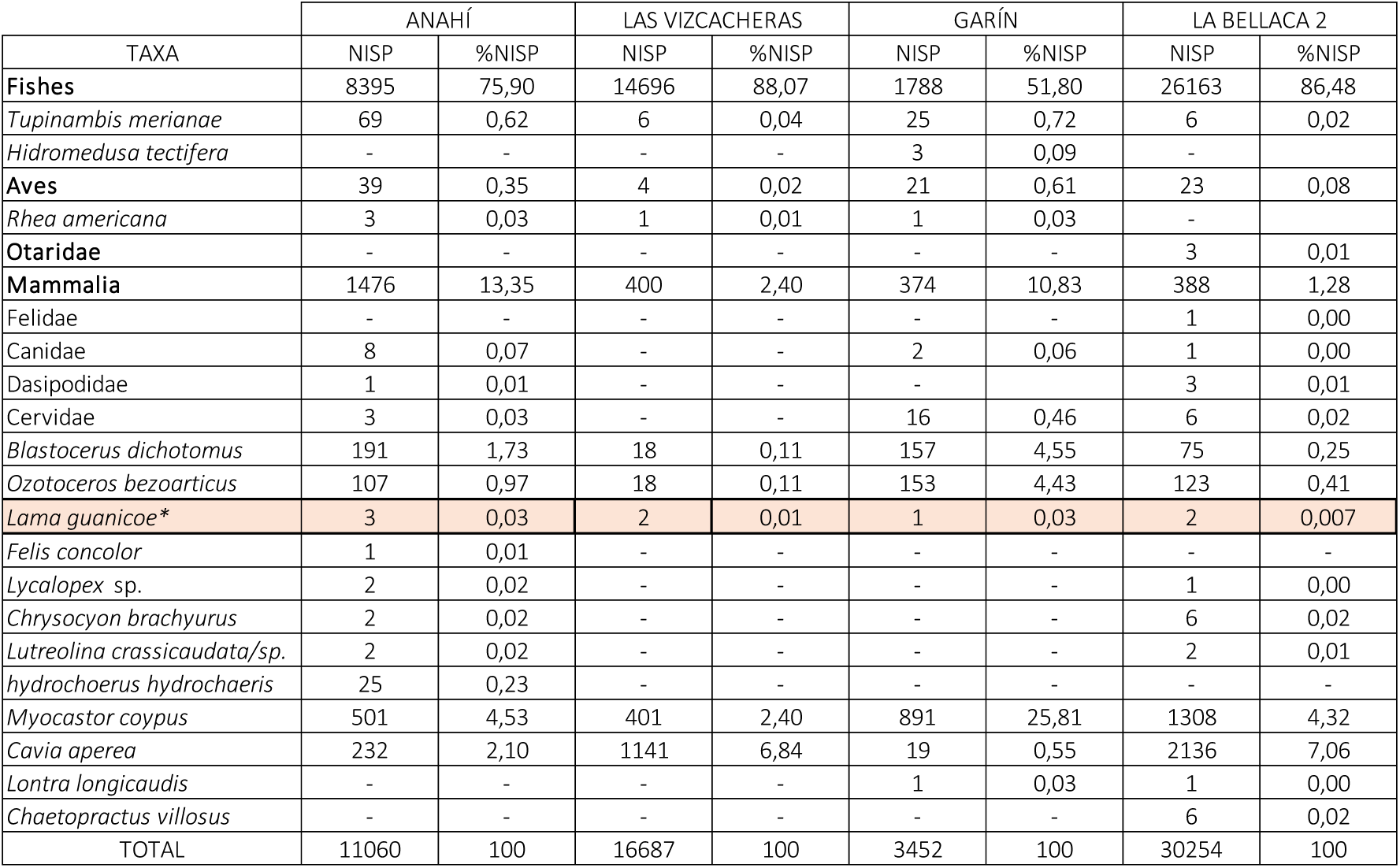
Faunal assemblages from some of the sites located at the wetland ecotone (taken and modified from Loponte, 2008). * The identification of camelid remains as *L. guanicoe* was initially made by Loponte (2008) due to their morphometric similarity, the absence of published archaeological records of domesticated camelids in the adjacent Pampas plain, and the lack of reliable historical evidence of llamas in the region. According to the results of this study, these identifications are confirmed as accurate (see main text). The NISP value for *L. guanicoe* at the La Bellaca 2 site was originally based on a single identified remain, but further analysis of the collection allowed for the identification of a second bone element attributed to this taxon.

Plant resources were also abundant in the wetland, particularly in the ecotone with the Pampa plain, where a narrow strip of forest with numerous edible species developed (Loponte, 2008). Although 16^th^-century travelers described these groups primarily as hunters and fishers, some accounts mention that they also manipulated or cultivated certain plants, such as maize.

Approximately 30 archaeological sites have been excavated in the wetland ecotone, 21 of which are detailed in Supplementary Table 2. Among these, sites such as El Cazador 3, El Espinillo, Garín, La Bellaca 3, Médanos de Escobar, and Rancho Largo feature excavation areas exceeding 50 m², with some extending up to 100 m². The remaining sites have excavation areas of around 20 m², providing a robust and representative sample of the local archaeological record. Fine mesh sieving was used at all sites to ensure the recovery of small fish bones, which were found in large quantities (see Table 1). These sites represent small base camps where a variety of activities took place, including burials, pottery production and use, lithic and bone tool manufacturing, and the processing of hides, wood, and plants. A summary of this record, though somewhat outdated, is available in English in Loponte et al. (2006).

Despite the extensive excavations, camelid bones were consistently found in very low quantities compared to other faunal remains, never exceeding 0.1% of the total NISP. As an example, Table 1 provides the faunal list from four sites examined in this study, which adequately illustrates the rarity of camelid remains in the wetland ecotone sites. It is worth noting that the camelid remains at these sites are intermixed with the rest of the fauna, showing no evidence of association with specific activities (religious or symbolic). They do not display any particular distribution within the record, nor do they exhibit different preservation states. These bones are predominantly phalanges and the distal ends of metapodials, with only a few exceptions listed in Supplementary Table 2. The same occurs with rhea bones (Acosta et al. 2023). Therefore, the peculiar representation of camelids at these sites cannot be attributed to sampling biases or differential preservation.

The northern Pampean plain adjacent to the Paraná River valley wetland is a completely different environment, as it is a grassland plain with almost no trees and very little surface water, mainly concentrated in small rivers and streams. Until the 16^th^ century, large herds of ungulates such as the Pampas deer (*O. bezoarticus*) and guanaco (*L. guanicoe*), as well asl rheas (*R. americana*), and some small mammals like the vizcacha (*Lagostomus maximus*) and armadillos (Dasypodidae) were found in this region. The Late Holocene (< 3.0 ky BP) archaeological sites in this plain, though much less excavated than those in the wetland, have relatively simple records. The faunal assemblages are dominated by guanaco bones (∼ >80% NISP), followed by Pampa deer and the aforementioned small mammals, with scarce remains of rhea, which was a difficult prey to capture (Acosta et al., 2023). The bone assemblages are generally well-preserved, with weathering stages primarily ranging between 1 and 3 on the Behrensmeyer (1978) scale. These sites contain little or no pottery but have a moderate number of lithic artifacts made from rocks sourced from the southern hills of this vast plain. These assemblages reflect highly mobile hunter-gatherer bands with simple technology with a subsistence based on guanaco.

Sixteenth-century historical sources describe the presence of small, highly mobile human groups that followed the movements of game across the interior of this vast plain, which multiple historical sources depict as sparsely populated. A key historical reference for our study, first noted by Loponte (1996-1998), indicates that these hunter-gatherer bands traded ‘sheep’ skins with indigenous groups settled in the adjacent wetland. The term ‘sheep’ was frequently used by the Spanish to refer to South American camelids, as they had no closer European equivalent than sheep, although they occasionally referred to them as ‘camels’ (Loponte & de Santis, 1995). Unfortunately, there is no English summary of the archaeology of this area yet, as this is a work in progress, but some studies in Spanish can be found in Ameghino (1947), Loponte (1996-1998; 2008), and Loponte et al. (2010).

### 2. The use of osteometry to differentiate camelid species

The osteometric differentiation between llamas and guanacos presents several uncertainties. Some bones, particularly those displaying greater robustness or size, as well as certain pathologies like exostosis, have been tentatively linked to stress from human management of domesticated animals (Yacobaccio, 2021). However, these indicators remain highly uncertain due to equifinality and are difficult to identify in archaeological samplings. Currently, the focus on distinguishing between llamas and guanacos based on osteometry centers on the dimensions of the first phalanges, but there is ongoing debate about the level of confidence in this method (e. g., Cartajena et al., 2007; del Papa, 2020; Elkin, 1996; Gasco & Marsh, 2015; Grant, 2010; Le Nuen et al., 2023; L’Heureux, 2005; Medina et al., 2014; Mondini & Muñoz, 2017; Yacobaccio, 2010, 2021, among others).

The second major issue is the dimensional overlap between llama and guanaco phalanges. Some authors (e.g., Gasco & Marsh, 2015; Yacobaccio, 2021) have suggested that the larger phalanges, which are the only ones that do not overlap with those of guanacos, likely belong to llamas. However, these authors also highlighted a key complication when comparing guanaco sizes, due to their clinal variation, with southern guanacos (from the Pampa and Patagonia) being larger. This size increase results in greater overlap in measurements between guanacos and llamas (Gasco & Marsh, 2015; Le Nuen et al., 2023). Consequently, the phalanges of Patagonian and Pampean guanacos overlap more with those of llamas, making these measurements less reliable for species identification in non-Andean regions. Further complicating the situation are hybrid populations (Cartajena et al., 2007) and past guanaco populations for which we lack adequate osteometric records. Together, these factors, along with new osteometric analyses cast doubt on the reliability of morphometrical criteria (Le Nuen et al., 2023).

From a dimensional perspective, it is at least possible to assess whether the first phalanges of guanacos from the Pampean plain and those from the ecotone share similar dimensions. Given that our samples vary in integrity due to fractures and that camelid remains are extremely scarce at the wetland sites, we compared a small sample of phalanges from sites located in both the northern Pampean plain and the wetland (Table 2). The results show no significant differences in measurements, suggesting that adult specimens from both regions were similar in size. However, as previously mentioned, the osteometric approach does not resolve the taxonomic ambiguity in our case study, so we present this data as supplementary information.

**Table 2.** Means and confidence intervals of the dimensions obtained from the first phalanx of guanacos recovered from the wetland and Pampa plain sites, expressed in mm (Gl, Bp -Bfp-, and Bd after von Den Driesch, 1976; FP1V3 and FP1V5, after Kent 1982) (taken from Buc & Loponte, 2016)

| Wetland samples |  |  |  |  | Interval of confidence (95%) |  |
| --- | --- | --- | --- | --- | --- | --- |
| Variable | Parameter | Estimate | S.E | *n* | Lower Limit | Upper limit |
| Gl | Mean | 77.23 | 1.11 | 6 | 73.66 | 80.09 |
| Bfp | Mean | 20.13 | 1.03 | 6 | 17.49 | 22.78 |
| FP1V3 | Mean | 19.04 | 0.47 | 6 | 17.84 | 20.25 |
| Bd | Mean | 16.83 | 0.32 | 5 | 15.94 | 17.72 |
| FP1V5 | Mean | 18.92 | 1.39 | 5 | 15.08 | 22.77 |

| Pampa plain samples |  |  |  |  | Interval of confidence (95%) |  |
| --- | --- | --- | --- | --- | --- | --- |
| Variable | Parameter | Estimate | S.E | *n* | Lower Limit | Upper limit |
| Gl | Mean | 75.50 | 1.38 | 6 | 71.96 | 79.04 |
| Bfp | Mean | 20.33 | 1.33 | 3 | 14.62 | 26.05 |
| FP1V3 | Mean | 18.93 | 0.43 | 4 | 15.55 | 20.30 |
| Bd | Mean | 16.63 | 0.39 | 5 | 15.55 | 17.71 |
| FP1V5 | Mean | 17.88 | 1.18 | 5 | 14.60 | 21.16 |

### 3. Key environmental indicators for this study

#### 3.1. δ^18^O regional values

The Paraná River serves as the primary water source for the Lower Paraná wetland. This river is extraregional, with its headwaters located in southern and central Brazil, where it captures rainfall from the Intertropical Convergence Zone. The meteoric water that precipitates in the Paraná River’s headwaters undergoes several fractionation episodes before reaching the wetland. The δ^18^O values of CO_3_ in bone apatite and biogenic carbonates from local organisms in the Lower Paraná wetland, including the ecotone adjacent to the Pampas plain, range between −5.1 and −2.4 ‰ V-PDB (Loponte & Ottalagano, 2022) (see Figure 1). This broad range results from various Dansgaard effects, such as the “amount effect” of precipitation, which fluctuates with hydrological pulses, as well as water evaporation in local lakes and streams obstructed by sediments and vegetation. In the neighbouring Pampas plain, biologically available water primarily comes from intermittent local rainfall and water stored in vegetation. Evaporation processes are more pronounced here, leading to slightly higher δ^18^O values in biogenic apatite, which range from −2.1 to 1.6 ‰ (Loponte & Ottalagano, 2022). During the late Holocene (after 3,000 years BP), both the wetland and the Pampas plain were influenced by the Medieval Climatic Anomaly (ca. 950–1250 CE) and the Little Ice Age (ca. 1450– 1880 CE). Despite these climatic fluctuations, no substantial changes in δ18O values have been observed (Loponte & Corriale, 2019; Loponte & Ottalagano, 2002).

**Figure 1.**
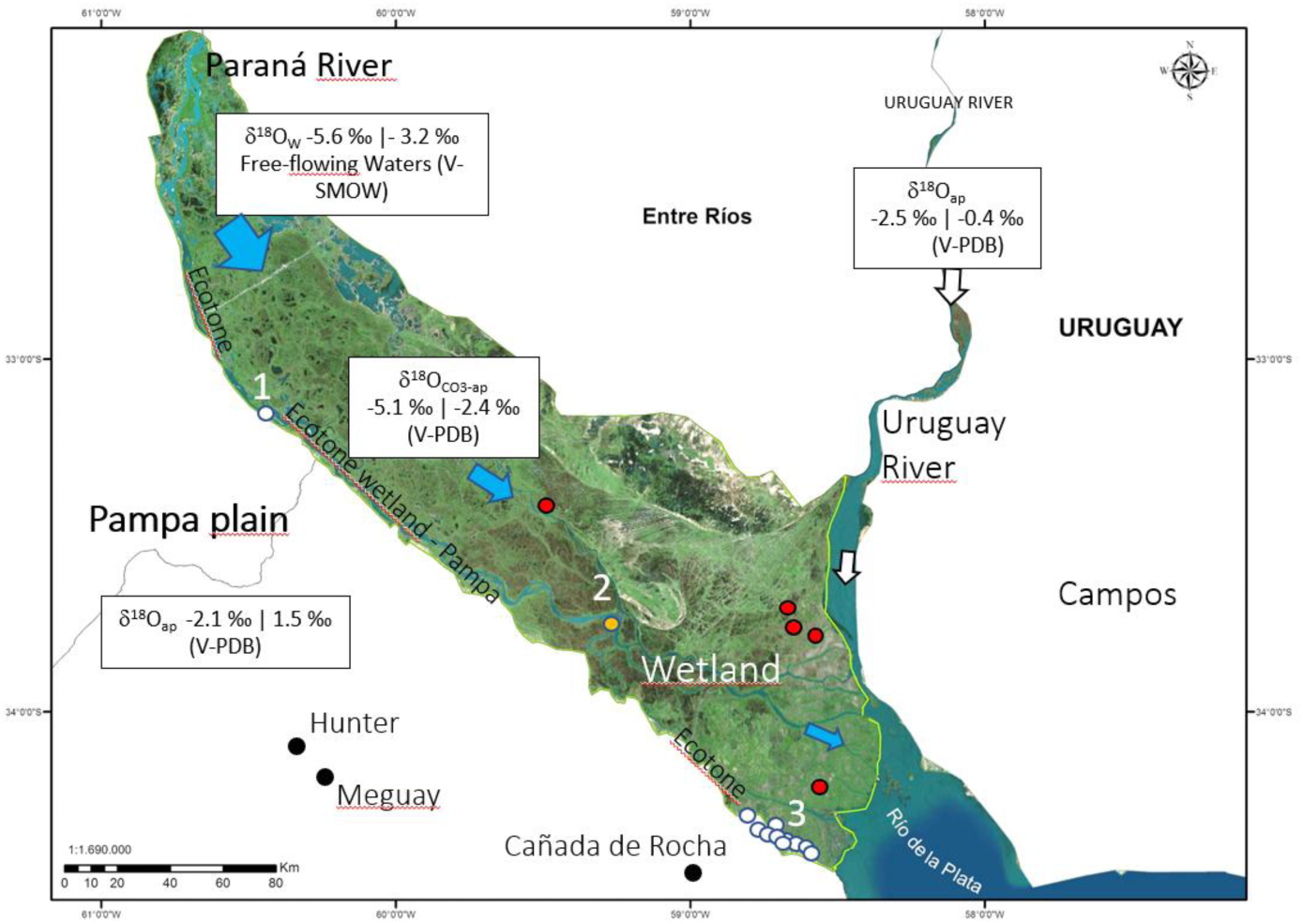
δ^18^O values of the region in local organism (bioapatite and biogenic carbonates and free-flowing water). The white circles represent the sites in the ecotone with the Pampas plains (or near to in the case of the orange dot). The red dots correspond to sites deep inside in the wetlands Taken and modified from Loponte & Ottalagano (2022).

The δ^18^O values of the Uruguay River, another major river bordering the wetland, differ from those of the Paraná due to its proximity to the South American Convergence Zone, where rainfall undergoes fewer fractionation events before reaching the river’s headwaters. However, the Uruguay River is not relevant to this study, as it is limited to the eastern sector of the wetland, which is outside the scope of our analysis.

#### 3.2. Geology of the area

The surface geology of the Pampas plain adjacent to the wetland corresponds to the Pampean Formation, characterized by aeolian loess deposits from the Upper Pleistocene. These deposits consist of massive brown to reddish silts that uniformly blanket the plain (Figure 2). In contrast, the surface geology of the Lower Paraná wetland is far more complex due to a middle Holocene marine ingression and the influence of marine tides throughout the Holocene. These tides transported sediments from the south to the north, which were subsequently reworked by fluvial processes that moved silts, clays, and sands southward (Figure 3). As a result, the wetland is characterized by the intersection of marine sands and coquinas with estuarine and fluvial sediments, creating a geomorphologically diverse and patchy environment (Figure 3).

**Figure 2.**
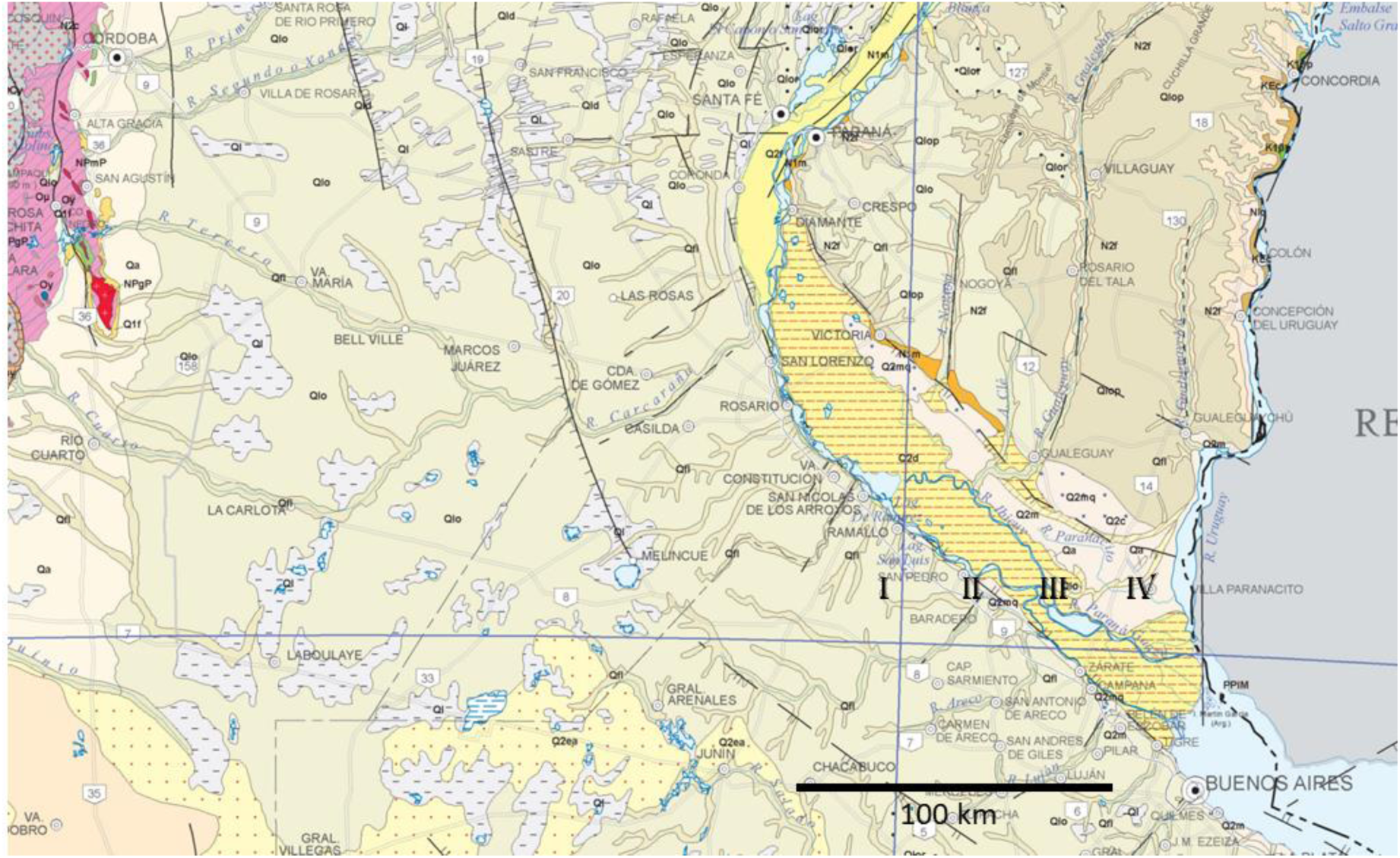
Geomorphological units of the studied region. I = Upper Pleistocene loess deposits of the Pampas plain. II = Pampas plain - wetland ecotone. III – IV: Sedimentary complex (marine and fluvial sediments) of the Lower Paraná wetland. Map taken and modified from Segemar (1997).

**Figure 3.**
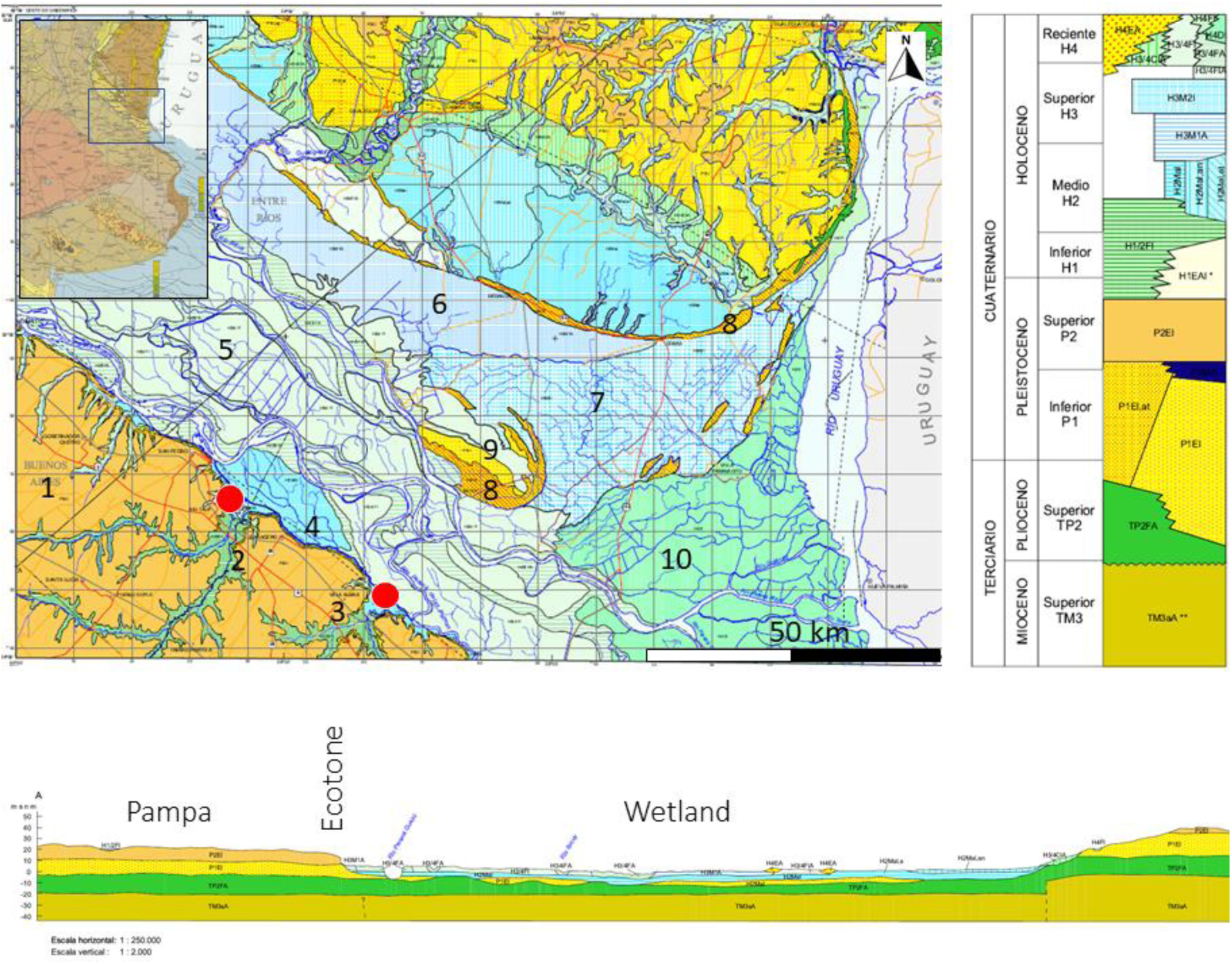
Geological map of the lower Paraná and adjacent areas. The red dots are the location of the sites in the ecotone, enclosed within circles of ⁓ 10 km diameter. 1 = Upper Pleistocene loess deposits. 2 = Recent Holocene deposits (< 3000 BP) of brown and green clays. 3 = Early Holocene fluvial deposits of reddish and greenish clays, with cross-bedded sands. 4 = Mid-Holocene tidal flat deposits with grayish-brown clays and silts. 5 = Late Holocene fluvial deposits with cross-bedded sands and brown silts within interdistributary plains. 6 = Mid to Late Holocene marine shoreline ridge deposits, with sandy beaches and shell beds. 7 = Estuarine shoreline ridge deposits with brown clays and sands. 8 = Active sand dunes. 9 = Pleistocene loess deposits. 10 = Deltaic plain deposits, with brown and gray sands and clays from the late Holocene to recent ages. Map taken and modified from Pereyra et al. (2002).

At the ecotone between the wetland and the Pampas plain, this complex sedimentary system merges with the Pampean loess and various fluvial sediments carried by rivers from the plain into the ecotone. This convergence results in a highly diverse sedimentary environment along the ecotonal area (Figure 3). Additionally, deeper geological layers not represented in surface planimetry emerge within the ecotone, as seen in the stratigraphic profile of Figure 4. These layers are particularly evident in the cliffs separating the Pampas plain from the wetland and occasionally appear in estuaries of rivers descending from the plain, further contributing to the sedimentary variability of the ecotone.

**Figure 4.**
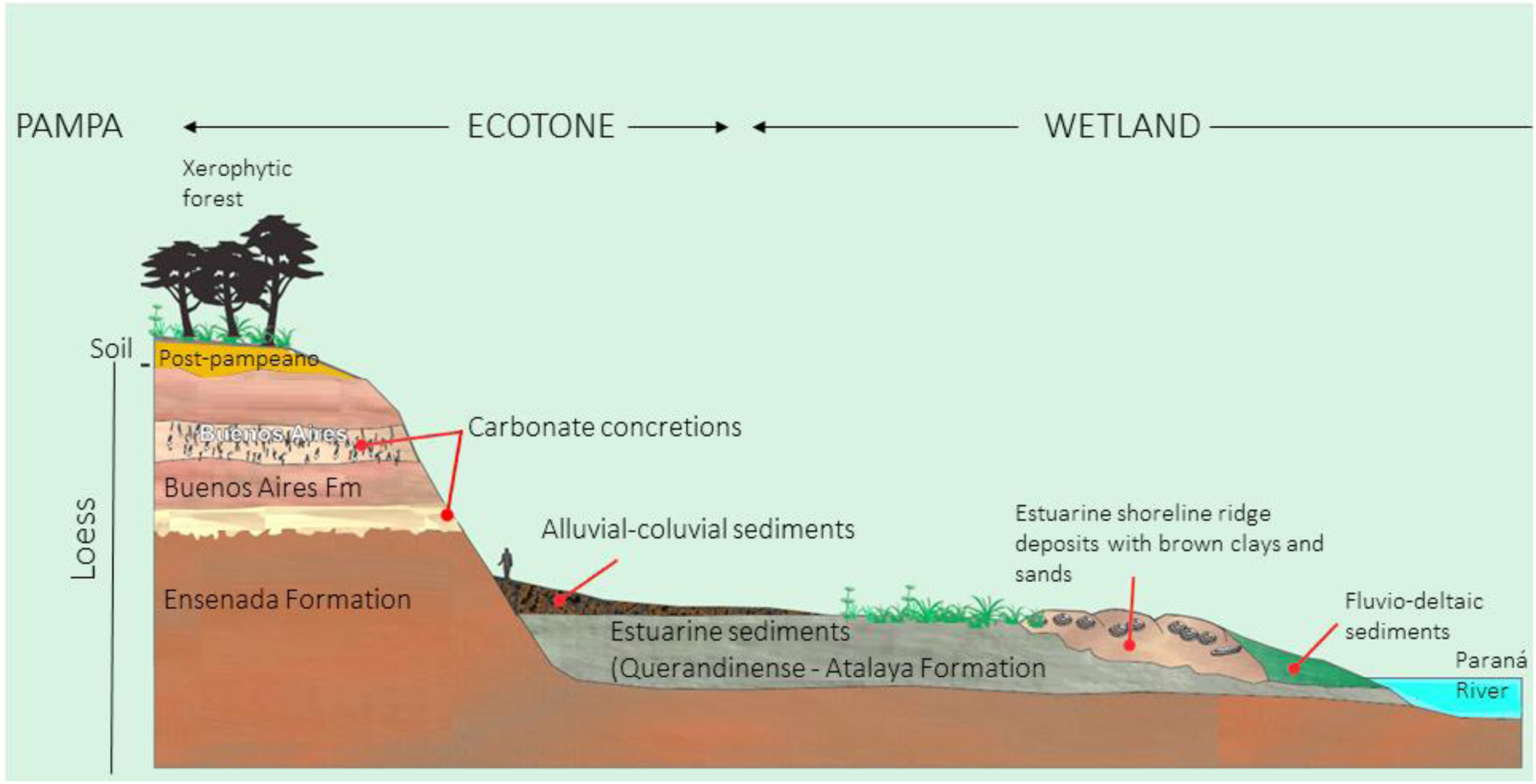
Cross-section of the study area, with a more detailed view of the ecotonal area. Image taken and modified from Miranda et al. (2020).

### 4. Isotopic analyses

δ^18^O and δ^13^C from CO_3_ were analyzed at the Environmental Isotope Laboratory, Dept. of Geosciences of the University of Arizona. Powdered samples were reacted with dehydrated phosphoric acid under vacuum at 70°C in the presence of silver foil. The samples were measured using an automated carbonate preparation device (KIEL-III) coupled to a gas-ratio mass spectrometer (Finnigan MAT 252). The isotope ratio measurement is calibrated based on repeated measurements of NBS-19 and NBS-18 and precision is ± 0.1 ‰ for δ^18^O and ±0.08‰ for δ^13^C (± 1σ). The apatite carbonate – CO_2_ oxygen isotope fractionation for the acid extraction is assumed to be identical to calcite. Removal of diagenetic carbonates follow the pre-treatment methods described in Koch et al. (1997). A second set of samples were analyzed at Instituto de Geocronología y Geología Isotópica (CONICET-Argentina) following a protocol similar to the previous one. The analysis of the isotopic composition of oxygen follows McCrea (1950) protocol, with some minor modifications.

The CO_3_ was converted into CO_2_ by reaction with phosphoric acid in a bath at 60 °C for two hours. The CO_2_ was purified in a vacuum line, using cryogenic traps to remove other volatile compounds (H_2_O vapor and eventually SO_2_) while the O_2_ and N_2_ were pumped since they are not condensable gases. The CO_2_ was analyzed in a Delta S Finnigan MAT mass spectrometer. The standard was the Carrara marble, which is calibrated with the NBS-19 and IAEA-CO-1 standard. The analytical error is ± 0.1 ‰. Interlaboratory bias between both sets of results are not significant for this study (Loponte & Ottalagano, 2022). All δ^18^O and δ^13^Cap values are expressed in per mill relative to VPDB.

For ^87^Sr/^86^Sr, analyses were carried out at the laboratories of the Centro Interdipartimentale Grandi Strumenti of the Università di Modena e Reggio Emilia (Italy). The samples were cleaned with MilliQ water and digested in concentrated HNO_3_. Strontium was separated with standard ion-exchange separation methods using the Eichrom Sr-spec resin. ^87^Sr/ ^86^Sr isotope ratios were measured by MC–ICP-MS (Thermo Scientific Neptune) using a bracketing sequence (blank-standard-blank-sample-sample-blank) to correct for possible instrumental drifts.

Repeated analyses of the NBS–987 standard provide an ^87^Sr/ ^86^Sr ratio of 0.710250 ± 0.000016 (2sd, N = 17). Total Sr blank did not exceed 100 pg. The ^87^Sr/^86^Sr ratios were normalized to ^86^Sr/^88^Sr ratio of 0.1194 and corrected for machine bias to an NBS–987 value of 0.710248 (Mc Arthur et al. 2001).

